# Long-range inhibitory axons from medial entorhinal cortex target lateral entorhinal neurons projecting to the hippocampal formation

**DOI:** 10.1101/2022.11.29.518323

**Authors:** Eirik S. Nilssen, Bente Jacobsen, Thanh P. Doan, Paulo J. B. Girão, Menno P. Witter

## Abstract

The functionally different lateral entorhinal cortex (LEC) and medial entorhinal cortex (MEC) are strongly interconnected. The role of this interconnectivity in view of their functional differences is not known. Here we provide details on a circuit that directly connects MEC to neurons in the superficial layers of LEC. Using a combination of anatomical tracing experiments and *in vitro* electrophysiological recordings in the mouse, we report that axons from MEC somatostatin-expressing GABAergic neurons densely distribute in layer I of LEC, where they drive strong and near selective inhibition of principal neurons in layer IIa. This inhibitory pathway is accompanied by MEC glutamatergic axons that innervate multiple layers of LEC and preferentially synapse onto principal neurons in layers IIb and III rather than principal neurons in layer IIa. These findings indicate that excitatory and inhibitory projections from MEC may separately regulate the activity of different populations of hippocampal-projecting principal neurons in LEC.

## INTRODUCTION

In the mammalian brain, the hippocampal formation (HF) and the parahippocampal region form a system that supports spatial navigation and episodic memory. A critical component of this system is the entorhinal cortex (EC), which provides most of the cortical input to the HF. I t is well-accepted that the rodent EC has two main subdivisions commonly referred to as the lateral EC (LEC) and the medial EC (MEC). These entorhinal areas have different functions and are thus believed to play complementary roles relevant to the formation of episodic memories (reviewed in e.g. Sugar and Moser 2019). There is strong evidence that MEC represents space through a manifold of functionally specialized neurons, such as grid cells (Fyhn et al., 2004; Hafting et al., 2005; Yartsev et al., 2011; Jacobs et al., 2013), object-vector cells (Høydal et al., 2019), border cells (Savelli et al., 2008; Solstad et al., 2008), speed cells (Kropff et al., 2015; Ye et al., 2018) and head-direction cells (Sargolini et al., 2006; Giocomo et al., 2014), whose collective activity is thought to map self-location as the animal navigates in space. In contrast, spatial coding is much less robust in LEC when animals explore open field environments (Hargreaves et al.,2005; Yoganarasimha et al., 2011). Instead, LEC is believed to primarily encode the content, e.g. object and odour-related information (Deshmukh and Knierim, 2011; Xu and Wilson, 2012; Tsao et al., 2013; Igarashi et al., 2014; Wang et al., 2018), as well as temporal sequences of ongoing experiences (Tsao et al., 2018; Bellmund et al., 2019; Montchal et al., 2019; Bright et al., 2020).

Although the spatially modulated cells characteristic of MEC are absent in LEC, the latter area has been shown to exhibit task-relevant processing of spatial information, often tied to the introduction of objects in the environment (Keene et al., 2016; Rodo et al., 2017). For example, some neurons in LEC develop spatially confined firing fields signaling the present or past location of objects (Deshmukh and Knierim, 2011; Tsao et al., 2013). Such spatial representations in LEC are possibly inherited from the HF or MEC, or both, because both areas project to LEC and are considered crucial elements of the viewpoint-independent (allocentric) spatial map in the medial temporal lobe. While hippocampal projections to LEC are known to preferentially target layer V (Rosene and Van Hoesen, 1977; Sürmeli et al., 2015; Ohara et al., 2018), axons deriving from neurons in layers II and V-VI of MEC densely innervate the superficial layers of LEC (Köhler,1986; Dolorfo and Amaral, 1998; Ohara et al., 2019), pointing to MEC as a likely candidate for relaying spatial information to neurons in the superficial layers of LEC. These superficial neurons, in turn, give rise to the lateral perforant path projecting to HF, suggesting that MEC may directly influence the activity of hippocampal-projecting neurons in LEC. In the pathway from LEC to the HF, hippocampal dentate gyrus (DG) and CA2/CA3 receive synaptic inputs from reelin-expressing principal neurons in layer IIa, whereas CA1 and subiculum are targeted by principal neurons in layer III as well as some calbindin-expressing neurons in layer IIb (van Groen et al., 2003; Kitamura et al., 2014; Kohara et al., 2014; Ohara et al., 2019). While it was recently shown that in the circuit linking LEC to MEC, neurons in layer IIa of LEC provide excitatory input, as well as elicit feed-forward inhibition, to excitatory neurons in layer II of MEC (Vandrey et al., 2022), the organizing principles that govern the reverse circuit connecting MEC to LEC have not been yet identified.

In this study, we used *in vitro* electrophysiology and neuroanatomical tracing to investigate the associative network connecting MEC to LEC, with a focus on neurons in LEC providing synaptic input to the HF. We find that this pathway is composed of excitatory and long-distance projecting inhibitory axons, and further present evidence that optogenetic activation of these projections has neuron-specific impact in superficial layers of LEC. Activation of somatostatin-expressing (SST^+^) GABAergic axons from MEC almost exclusively inhibit principal neurons in layer IIa (P_IIa_), whereas inputs from MEC glutamatergic axons preferentially excite principal neurons in layers IIb (PN_IIb_) and III (PN_III_). Our data provide insight into the organizing principles ruling the MEC–LEC circuit and indicate a role for MEC in exerting selective modulation of the activity of hippocampal-projecting neurons in LEC.

## RESULTS

### Glutamatergic neurons and somatostatin-expressing GABAergic neurons in MEC send axons to LEC

We first aimed to identify MEC neurons that send axons to LEC. Retrograde neuroanatomical tracing was conducted in GAD67*^eCFP^* mice, which provide genetic access to GABAergic neurons (Tamamaki et al., 2003), to distinguish between glutamatergic and potential GABAergic contributions to this pathway. Injections of the chemical tracer fluorogold (FG, n = 2 mice) or an adeno-associated virus (AAV2-CAG-tdTomato, n = 1 mouse) into dorsal LEC revealed that retrogradely tagged neurons in MEC were found in both deep and superficial layers (deep, V-VI; superficial, I-III; data from 3 GAD67*^eCFP^* mice; Fig. 1A and Supplementary Fig. 1). These results corroborate earlier reports that axons projecting to LEC originate from neurons in both deep and superficial layers of MEC (Köhler, 1986; Dolorfo and Amaral, 1998). Out of a total of 5190 counted FG/tdTomato+ neurons in MEC (n = 3 mice), 3.7 ± 2.4% (mean ± s.d) expressed the GFP-signal of GAD67-positive neurons. Since many corticocortical long-distance projecting GABAergic neurons, i.e., GABAergic neurons whose axons project outside of their anatomical area of origin, synthesize the neuropeptide somatostatin (Melzer and Monyer, 2020), we examined if SST^+^ neurons are found among the GABAergic neurons projecting from MEC to LEC. Immunostaining of the tissue revealed that 64.7 ± 22.6% (mean ± s.d) of retrogradely labelled GABAergic neurons were immunoreactive for SST (Figs. 1B and 1C).

**Figure 1.**
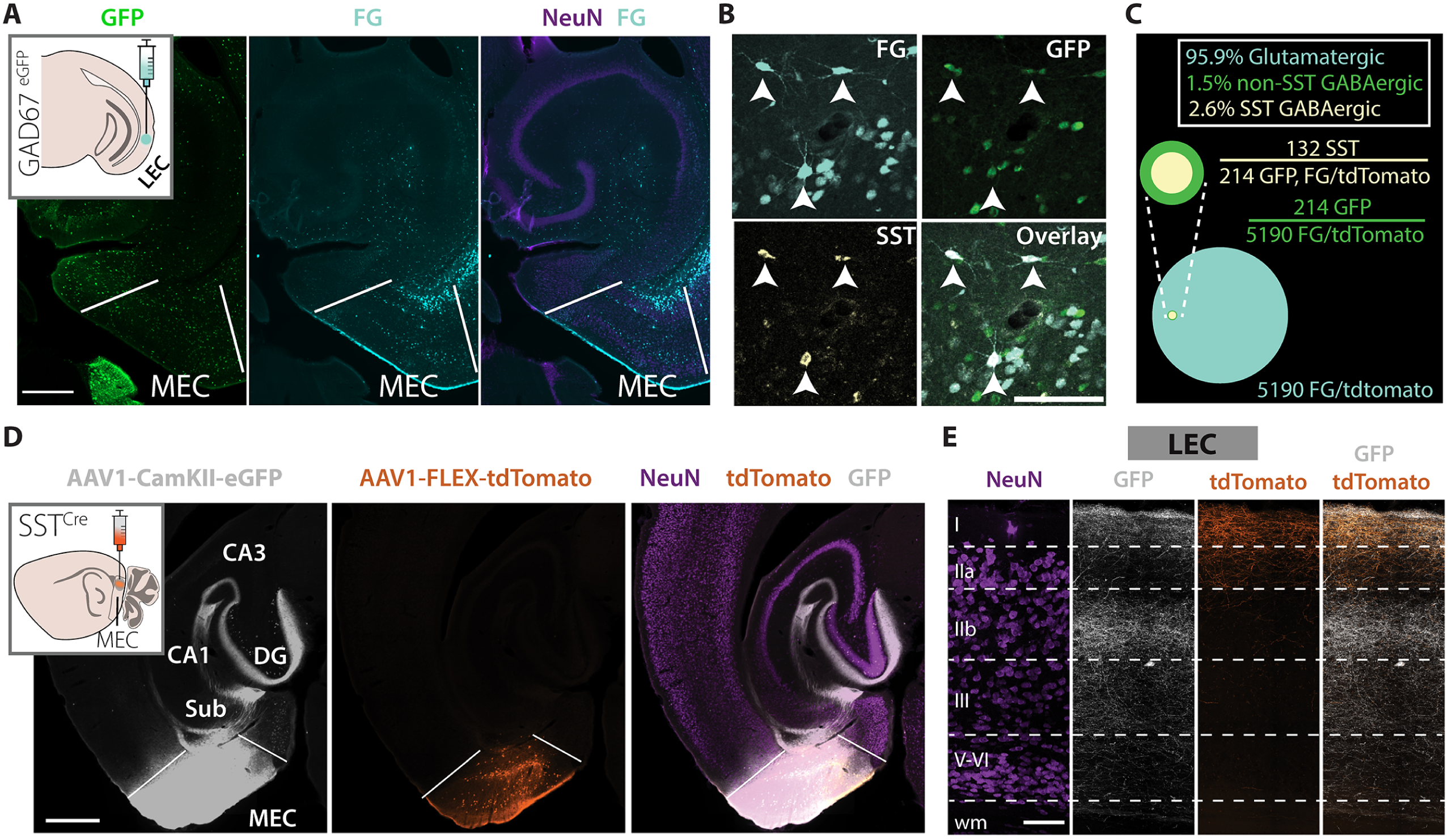
Glutamatergic neurons and SST^+^ GABAergic neurons project from MEC to LEC. **(A)** Horizontal brain section through MEC of a GAD67^eGFP^ mouse showing GFP^+^ neurons (green) together with retrogradelly labelled neurons (light cyan) from LEC. Inset: Schematic of the retrograde tracer injection in LEC. Scale bar, 500 µm. See Supplementary Fig. 1 for images showing the dorsoventral extent of the injection site. **(B)** SST^+^ GABAergic neurons in MEC project to LEC. Arrowheads indicate examples of triple-labelled neurons (SST^+^, GAD67*^eCFP^*, FG) from the experiment in **A**. Scale bar, 100 µm. **(C)** Proportions of GABAergic neurons (SST^+^ and non-SST^+^) and glutamatergic neurons in the MEC-LEC pathway (neuron counts from 3 mice). **(D)** Delivery of an equal mix of AAV1-CAG-FLEX-tdTomato and AAV1-CamKII-eGFP-WPRE-rBG into MEC of SST^Cre^ mice (upper left schematic) selectively labels SST^+^ neurons with tdTomato and glutamatergic neurons with GFP. Sub, subiculum. Scale bar, 500 µm. See Supplementary Fig. 2 for images showing the dorsoventral extent of the injection site. **(E)** Different laminar innervation patterns in LEC of glutamatergic axons and SST^+^ axons from the experiment in **D**. Wm, white matter. Scale bar, 100 µm.

We independently labelled SST neurons and glutamatergic neurons by injecting a mixture of two viruses into MEC of transgenic SST^Cre^ mice (n = 3) to investigate medial entorhinal innervation patterns in LEC. The virus solution had an equal ratio of i) an AAV vector driving GFP expression in glutamatergic neurons under the CamKII promoter (AAV1-CamKII -eGFP-WPRE-rBG) and ii) a Cre-dependent AAV vector encoding tdTomato (AAV1-CAG-FLEX-tdTomato). CamKII-promoted expression led to extensive axonal labelling in the HF and in LEC layers I, IIb and III, with additional weaker axonal labelling in layers IIa and V-VI. The Cre-driven fluorescence of SST^+^ neurons was distributed across layers of MEC, with an additional dense axonal labelling in ipsilateral LEC. Within LEC, tagged axons were found in layers I and IIa, with some scattered axons also in layers IIb and III (Figs. 1D, 1E and Supplementary Figs. 2 and 3A, B). Virally labelled SST axons were not observed elsewhere in the hippocampal-parahippocampal region. We carefully checked all sections at and close to the levels we used for our experiments and did not observe any virally labelled neurons in LEC.

In contrast to the dense innervation of LEC by medial entorhinal SST^+^ axons, only very few tagged axons were observed in MEC when targeting the LEC SST^+^ neuron population by injecting AAV1-CAG-FLEX-tdTomato into LEC (n = 2 SST^Cre^ mice; Supplementary Fig. 3C, D). Most cortical inhibitory neurons can be identified by their expression of either somatostatin, parvalbumin or vasoactive intestinal peptide, which are non-overlapping molecular markers used to classify main populations of GABAergic neurons (Tremblay et al., 2016). We therefore tested whether parvalbumin-expressing (PV^+^) or vasoactive intestinal peptide-expressing (VIP^+^) GABAergic neurons in MEC also supply long-range inhibition to LEC. In contrast to the clear innervation of LEC observed after labelling of MEC SST^+^ neurons (Fig. 1), we did not observe any labelled axons innervating LEC following conditional viral tagging of PV^+^ neurons or VIP^+^ neurons in PV^Cre^ (n = 5) or VIP^Cre^ (n = 3) mice, respectively (Supplementary Fig. 4). Thus, our data show that axons from MEC SST neurons, but not those of PV^+^ or VIP^+^ neurons, densely innervate the superficial layers of LEC. Although we have not examined whether all other molecularly defined inhibitory neurons, including those expressing the neuropeptide cholecystokinin, also send axons to LEC, our findings support that SST^+^ neurons are a main contributor to the inhibitory pathway from MEC to LEC. Additionally, this inhibitory neuron class is virtually absent from the reverse pathway that connects LEC to MEC.

To gain insight into the wider circuit that indirectly connects to LEC through the inhibitory MEC -LEC pathway, we mapped brain-wide monosynaptic inputs to MEC SST^+^ neurons via retrograde rabies virus tracing in transgenic SST^Cre^ mice. Hippocampal CA1 and subiculum contained more than 40 % of all counted neurons, while substantial numbers of presynaptic neurons were also found in presubiculum, medial septum, retrosplenial cortex and the anterodorsal thalamic nucleus (Supplementary Fig. 5). These findings point to brain areas supporting spatial navigation, i.e., the HF and the greater parahippocampal network, as important modulators of MEC SST^+^ neurons, likely including those SST^+^ neurons that project to LEC.

### Target preference of medial entorhinal axons in LEC

We next set out to map functional connections from MEC axons onto neurons in LEC using patch-clamp electrophysiology in acute brain slices. Injections of an AAV vector into MEC of wild-type mice enabled us to tag axons with the light-sensitive cation channel channelrhodopsin2 fused to a yellow fluorescent reporter protein (AAV1-hSyn-ChR2-YFP). Injection selectivity was verified by the presence of axonal labelling within the middle, but not the outer, third of the molecular layer of DG (Fig. 2A, left; Supplementary Fig. 6), a pattern consistent with hippocampal innervation by MEC (van Groen et al., 2003). Principal neurons in the superficial layers (layers IIa, IIb and III) of LEC were targeted for somatic whole-cell recordings. These layers are heavily innervated by medial entorhinal axons (Figs. 1E and 2A) and contain the main neuronal populations that project to the HF (van Groen et al., 2003; Kitamura et al., 2014; Ohara et al., 2019).

**Figure 2.**
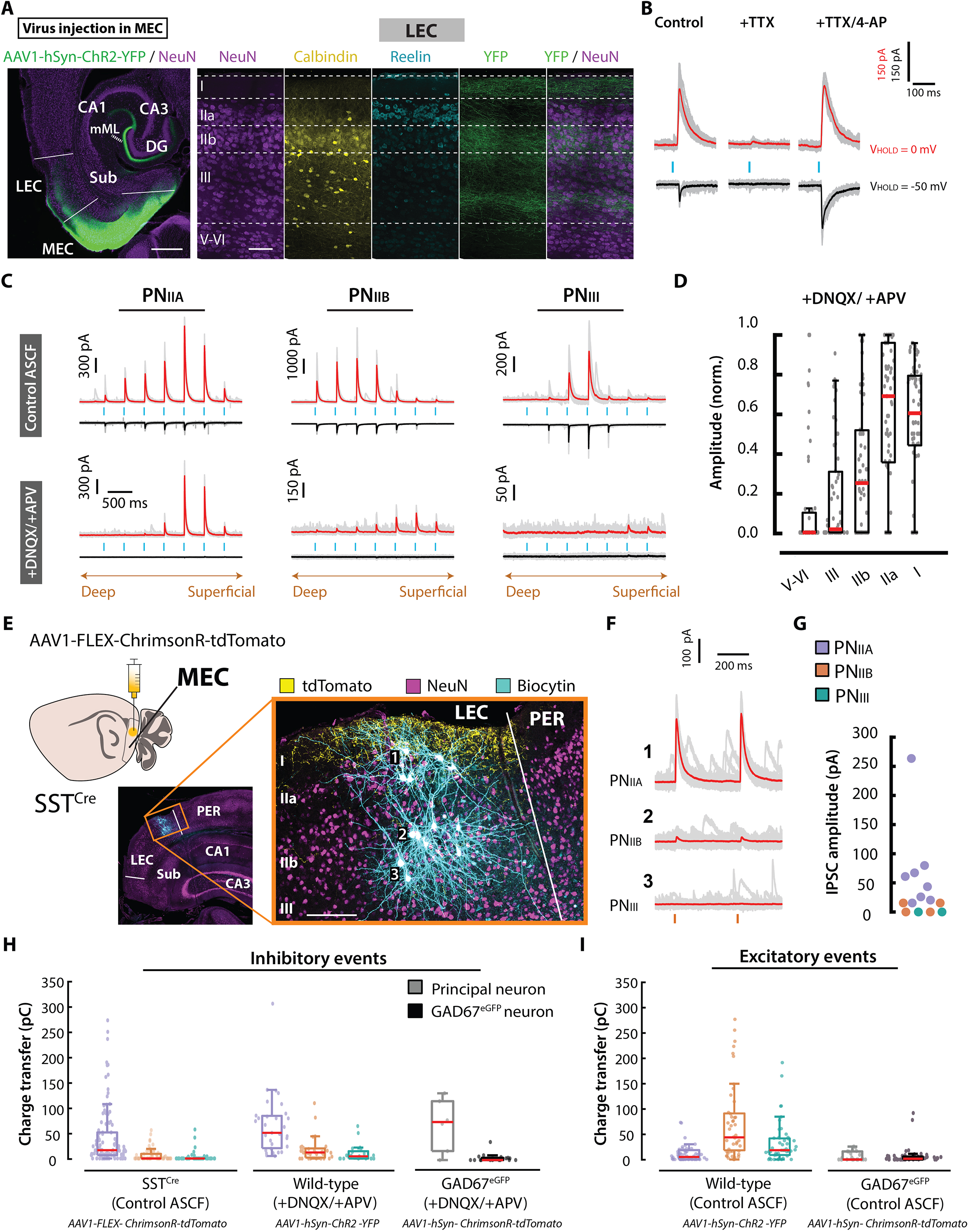
Target preference of medial entorhinal axons in LEC. **(A)** Left: Virus injection in MEC of a wild-type mouse. Right: High magnification image showing YFP-labelled photosensitive (ChR2) axons in LEC. MML, DG middle molecular layer. Scale bars, 500 µm (left) and 100 µm (right). **(B)** Membrane current recordings of a representative PN_IIA_ to the indicated pharmacological treatments during optogenetic stimulation (blue bars). The average membrane current traces (red/black) are superimposed on the individual traces (grey). **(C)** Laser-evoked EPSCs (black) and IPSCs (red) in representative PN_IIA_, PN_IIB_ and PN_III_ recorded without (top, control ACSF) or with glutamatergic synaptic blockers (bottom, DNQX/APV). **(D)** Normalized IPSC amplitudes plotted against the location of photostimulation. Boxes indicate 25th-75th percentiles, red line is median, whiskers extend to all data points not considered outliers. All data points are superimposed on the box plots. Each value represents data from one cell resulting from photostimulation of the given layer. **(E)** Patch-clamp recorded neurons together with photosensitive MEC SST^+^ axons in a semicoronal slice of LEC from an SST^Cre^ mouse. PER, perirhinal cortex. Schematic: Injection to virally label MEC SST^+^ neurons with ChrimsonR-tdTomato. Scale bars, 200 µm. **(F)** Membrane current recordings of the numbered neurons in **E** during photostimulation (orange bars). **(G)** Membrane current amplitudes recorded from all neurons in the experiment in **E**. **(H)** Charge transfers in response to optogenetic activation when clamping neurons at 0 mV. Non-responding neurons have charge transfer values of 0 pC. **(I)** Same as in **H**, but for data obtained when clamping neurons to −50 mV. The virus used in each experiment is indicated below each individual panel. The color code for PN_IIA_, PN_IIB_ and PN_III_ in **(H)** and **(I)** follow the color code as defined in **G**.

We recorded membrane currents while depolarizing photosensitive medial entorhinal axons by delivering brief laser pulses of blue light in discrete spots across layers of LEC. The membrane potential of recorded neurons was alternately clamped to the reversal potential for inhibitory and excitatory currents (Vinh = E*_cl_*_−_ ≈ −50 mV and V_exc_ ≈ 0 mV) to isolate excitatory and inhibitory synaptic events, respectively. Optogenetic mapping revealed laser-evoked excitatory postsynaptic currents (EPSCs) and inhibitory postsynaptic currents (IPSCs) in all three groups (PN_IIa_, PN_IIb_ and PN_III_; data from 14 mice). To distinguish between monosynaptic and polysynaptic responses, in a subset of experiments, we isolated monosynaptic currents by blocking voltage gated sodium channels with tetrodotoxin (TTX, 1 µM) and voltage gated potassium channels with 4-aminopyridine (4-AP, 100 µM; Petreanu et al. 2009). Some synaptic responses, both excitatory and inhibitory, were intact in PN_IIa_ (EPSC: n = 7neurons; IPSC: n = 12 neurons; 5 mice), PN_IIb_ (EPSC: n = 9 neurons; IPSC: 10 neurons; 4 mice) and PN_III_ (EPSC: n = 7 neurons; IPSC: 5 neurons; 3 mice), confirming that these three populations receive monosynaptic inputs from MEC (Fig. 2B). In agreement with the pattern of MEC glutamatergic and GABAergic axons, the pharmacological intervention revealed that monosynaptic excitatory inputs could be evoked from both deep and superficial layers, whereas monosynaptic inhibition was most prominent following photostimulation of superficial locations (layers IIb–I; Supplementary Fig. 7).

To further investigate the direct GABAergic input to LEC principal neurons, we isolated monosynaptic inhibitory responses by clamping neurons to the reversal potential for excitatory currents (≈0 mV) while blocking glutamatergic synaptic transmission (DNQX, 10 µM; APV, 50 µM). Under these conditions, there was a clear relationship between the amplitude of recorded inhibitory membrane currents and the location of photostimulation (p < 0.0001, Kruskal–Wallis test). Photostimulation of axons within layers I and IIa produced outward currents with amplitudes more than twice as large as those elicited by photostimulation of the other layers (n =66 neurons from 8 mice, pooled data from PN_IIa_, PN_IIb_ and PN_III_; p < 0.01, Kruskal–Wallis multiple comparison test; Figs. 2C and 2D). These were GABA_A_-mediated chloride currents as they were blocked by the GABA_A_ - antagonist bicuculline (10 µM, n = 11 neurons from 2 mice; Supplementary Fig. 8).

Because MEC SST^+^ axons densely innervate layers I/I Ia of LEC (Fig. 1E), we sought to directly test whether SST^+^ neurons are the source of the observed monosynaptic inhibition. AAV1-FLEX-ChrimsonR-tdTomato was injected into MEC of SST^Cre^ mice (n = 13) to conditionally express the red-shifted channelrhodopsin ChrimsonR in SST^+^ neurons. This yielded a dense plexus of photosensitive ChrimsonR^+^ axons in layer I of LEC (Fig. 2E). Optogenetic axonal activation readily produced outward synaptic currents in PN_IIa_, but such responses were weaker and less frequently observed among PN_IIb_ and PN_II_ (Figs. 2F and 2G). To determine whether MEC expresses a target preference for principal neurons in LEC, we analyzed the strength of the synaptic responses by measuring membrane charge transfers evoked by stimulation of photosensitive axons. First, the analyses were restricted to outward currents recorded when the membrane potential was clamped at 0 mV (Fig. 2H). In SST^Cre^ mice, larger charge transfers were found in PN_IIa_ than in PN_IIb_ or PN_III_ (p < 0.0001, Kruskal-Wallis multiple comparison test; n = 106 PN_IIA_, 54 PN_IIB_ and 50 PN_III_). Similarly, stronger responses were detected among PN_IIa_ compared with PN_IIb_ and PN_III_ when monosynaptic inhibitory inputs were pharmacologically isolated in wild-type mice (+DNQX/+APV; p<0.001, Kruskal-Wallis multiple comparison test; n = 31 PN_IIA_, 32 PN_IIB_ and 21 PN_III_; Fig. 2H). Second, when analyses were limited to laser-evoked excitatory events recorded by clamping neurons at −50 mV in wild-type mice, we observed charge transfers with the largest magnitude in PN_IIb_, followed by PN_III_ and PN_IIa_ (p < 0.05, Kruskal-Wallis multiple comparison test; n = 48 PN_IIA_, 45 PN_IIB_ and 45 PN_III_; Fig. 2I).

Lastly, we investigated if GABAergic neurons in LEC are additional recipients of synaptic inputs from MEC. We recorded from GFP GABAergic neurons (n = 51) and principal neurons (17 PN_IIa_, 2 PN_IIb_ and 4 PN_III_) across superficial layers of LEC in GAD67*^eCFP^* animals (n = 7) that were injected with AAV1-hSyn-ChrimsonR-tdTomato into MEC (Supplementary Fig. 9). By measuring the average laser-evoked charge transfer, we found that GABAergic neurons had comparably strong excitatory responses to principal neurons (p = 0.82, Wilcoxon rank sum test; Fig. 2I and Supplementary Fig. 9). In contrast, the median amplitude of the monosynaptic IPSCs recorded from GABAergic neurons was much smaller than that of principal neurons (+DNQX/+APV; n = 18 tested GABAergic neurons; n = 6 tested principal neurons; p = 0.0056, Wilcoxon rank sum test; Fig. 2H and Supplementary Fig. 9). Together, these data indicate that PN_IIa_ receive the strongest inhibition mediated by long-distance SST projections from MEC.

### Target-specific control of postsynaptic neuron activity by medial entorhinal axons in layer I

As layers I/IIa are innervated by axons from both GABAergic and glutamatergic MEC neurons (Fig. 1E), we asked how a combined output of axons in layer I impacts the activity of postsynaptic neurons. In slices prepared from 8 wild-type mice injected with AAV1-hSyn-ChR2-YFP into MEC, we carried out membrane potential recordings of PN_IIa_, PN_IIb_ and PN_III_ while optogenetically activating axons in layer I. Recorded neurons were experimentally depolarized to suprathreshold potentials to ensure an adequate driving force for both inhibitory and excitatory synaptic currents (Figs. 3A and 3B).

**Figure 3.**
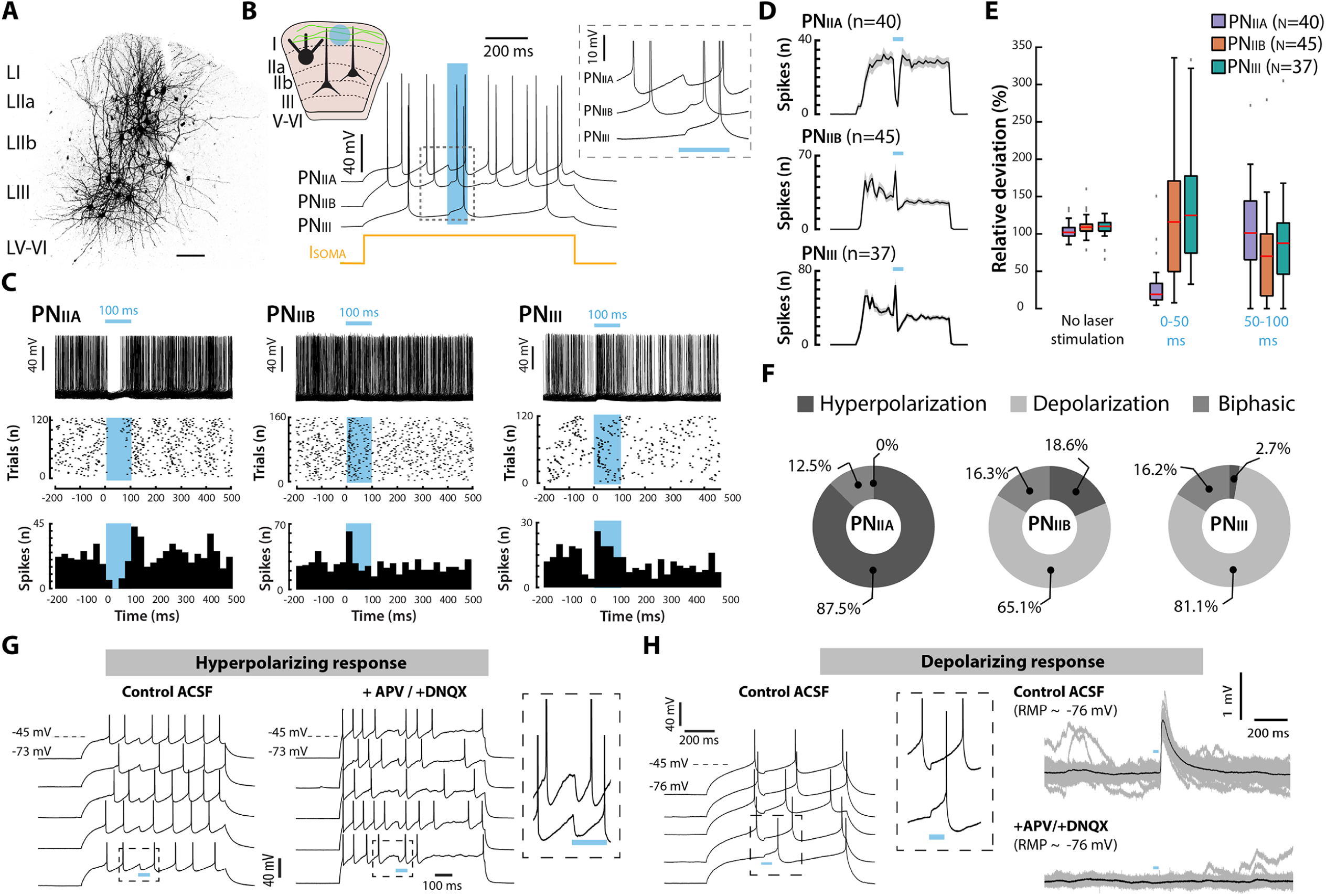
Target-specific control of postsynaptic neuron activity by medial entorhinal axons in layer I. **(A)** Biocytin-filled principal neurons recorded from layers IIa, IIb and III. Scale bar, 100 µm. **(B)** Different laser-evoked responses of PN_IIA_ versus PN_IIB_ and PN_III_ (PN_IIA_, hyperpolarizing; PN_IIB_, PN_III_; depolarizing). Neurons were depolarized by somatic current injections (yellow) while MEC axons were photostimulated with light (blue bar). Schematic upper left: A 100 ms laser stimulation (blue) of MEC axons (ChR2, green) in LEC layer I during current-clamp recordings. **(C)** Action potential firing across multiple sweeps of the photostimulation (blue bar) protocol in representative neurons. Top: Voltage recordings. Middle: Raster plots. Bottom: Histograms. Note the transient suppression of spiking activity in PN_IIA_ and the increase in spiking activity in PN_IIB_ and PN_III_ during photostimulation. **(D)** Mean firing rate (mean number of action potentials per bin) of all tested PN_IIA_, PN_IIB_ and PN_III_. Plots are mean (black) ± standard error of the mean (grey). **(E)** Box plots showing the deviation in action potential firing relative to the average firing activity without laser stimulation. Box plots as in **Fig. 2**. Outliers are shown as grey points. **(F)** Donut plots showing the proportion of laser-evoked responses. **(G)** Laser-evoked hyperpolarizing responses are resistant to application of glutamatergic synaptic blockers (DNQX/APV). The recording is from a representative PN_IIA_. **(H)** Left: Laser-evoked depolarizing responses recorded from a representative PN_III_. Right: The responses disappear when blocking glutamatergic synaptic transmission with APV and DNQX. RMP, resting membrane potential. In the rightmost panels the average voltage trace (black) is overlaid the individual voltage traces (grey).

The most striking difference in optogenetically induced responses was evident for PN_IIa_ versus PN_IIb_ / PN_III_, on the population level (Fig. 3) and between single neurons recorded in the same slice (Supplementary Fig. 10). Visual inspection of action potential firing revealed that PN_IIa_, unlike PN_IIb_ and PN_III_, fired very few spikes during blue light illumination (Figs. 3C and 3D). The median number of action potentials emitted by PN_IIa_ decreased to only 19 % of baseline levels (average number of action potentials without laser stimulation) during the first half (0–50 ms) of laser stimulation (n = 40, p < 0.0001). The median firing activity of PN_IIb_ and PN_II_ was not different from baseline, but the variability in the number of emitted spikes was high (p = 0.26; Wilcoxon signed rank test; Fig. 3E). During the second half (50–100 ms) of laser stimulation, the spiking activity of PN_IIa_ recovered while the firing activity of both PN_IIb_ and PN_III_ decreased, the latter two showing a median action potential firing of 70% and 87 %, respectively, relative to baseline (PN_IIa_: p = 0.54, PN_IIb_: p < 0.0001, PN_III_: p = 0.02; Wilcoxon signed rank test; Fig. 3E).

The difference in spiking activity between PN_IIa_ and PN_IIb_/PN_III_ matched a clear difference in laser-evoked postsynaptic potentials. Responding PN_IIa_ invariably showed hyperpolarizing deflections of the membrane potential, which in around 10% of PN_IIa_ were accompanied by depolarizing deflections (biphasic response; n = 5/40 neurons). Exclusive depolarizing responses were not observed in this neuron group (Figs. 3B and 3F). In contrast, 65% of responding PN_IIb_ and 81% of responding PN_II_ showed pure depolarizing membrane potential deflections (PN_IIb_, n = 28/43; PN_III_, n = 30/37). Some responding PN_IIb_ and PN_III_ showed biphasic responses or solely hyperpolarizing deflections (PN _IIb_: biphasic, n = 7/43; hyperpolarizing, n = 8/43; PN_III_: biphasic, n = 6/37; hyperpolarizing, n = 1/37; Figs. 3B and 3F).

To assess whether the observed hyperpolarizing responses could reflect local inhibition driven by excitatory MEC inputs, we silenced any contributions from polysynaptic excitatory pathways by eliminating glutamatergic synaptic activity in the slice. Tested laser-evoked hyperpolarizing responses were intact under these conditions (n = 16 neurons from 3 mice), whereas tested depolarizing responses (n = 7 neurons from 3 mice) disappeared (Figs. 3G and 3H). First, these data confirm that hyperpolarizing responses are monosynaptic, which is consistent with synaptic inputs from photosensitive long-range medial entorhinal inhibitory axons rather than recruitment of feed-forward inhibitory mechanisms. Second, they show that depolarizing responses are mediated by synaptic inputs from a glutamatergic pathway. Taken together, co-activation of MEC glutamatergic and GABAergic axons in LEC layer I induces a net inhibitory drive to PN_IIa_ and a net excitatory drive to PN_IIb_ or PN_II_. In our hands, driving input from MEC axons in layer I therefore produces a selective inhibition of PN_IIa_ and a selective excitation of PN_IIb_ and PN_III_.

### Cortical glutamatergic inputs to principal neurons in layer IIa

In a separate set of experiments, we investigated whether inhibitory synaptic responses in PN_IIa_ could also be evoked by activating other cortical areas innervating the superficial layers of LEC. In wild-type mice (n = 3), we first mapped the projection pattern in LEC of three regions previously reported to innervate LEC: the piriform cortex (PIR; Haberly and Price 1978; Kerr et al. 2007), the contralateral LEC (cLEC; Blackstad 1956; Leitner et al. 2016) and the perirhinal cortex (PER; Burwell and Amaral 1998; Doan et al. 2019). We confirmed that these areas send axons to the superficial layers of LEC using anterograde tracing. Axons from PIR and cLEC were confined almost exclusively to layers I and IIa whereas those from PER terminated diffusely within layers I– III (Fig. 4A and Supplementary Fig. 11). By retrogradely tracing these pathways in 3 GAD67*^eCFP^* mice, we observed that 8.6 ± 5.6% (mean ± s.d; 392 / 4651 FG/tdTomato^+^) of labelled neurons in PER and 0.4% ± 0.62 (mean ± s.d; 12 / 1853 FG/tdTomato^+^) of labelled neurons in PIR overlapped with GFP and thus were GABAergic. We did not detect any GFP GABAergic neurons within the projection from cLEC (518 FG/tdTomato^+^ neurons were counted; Fig. 4B).

**Figure 4.**
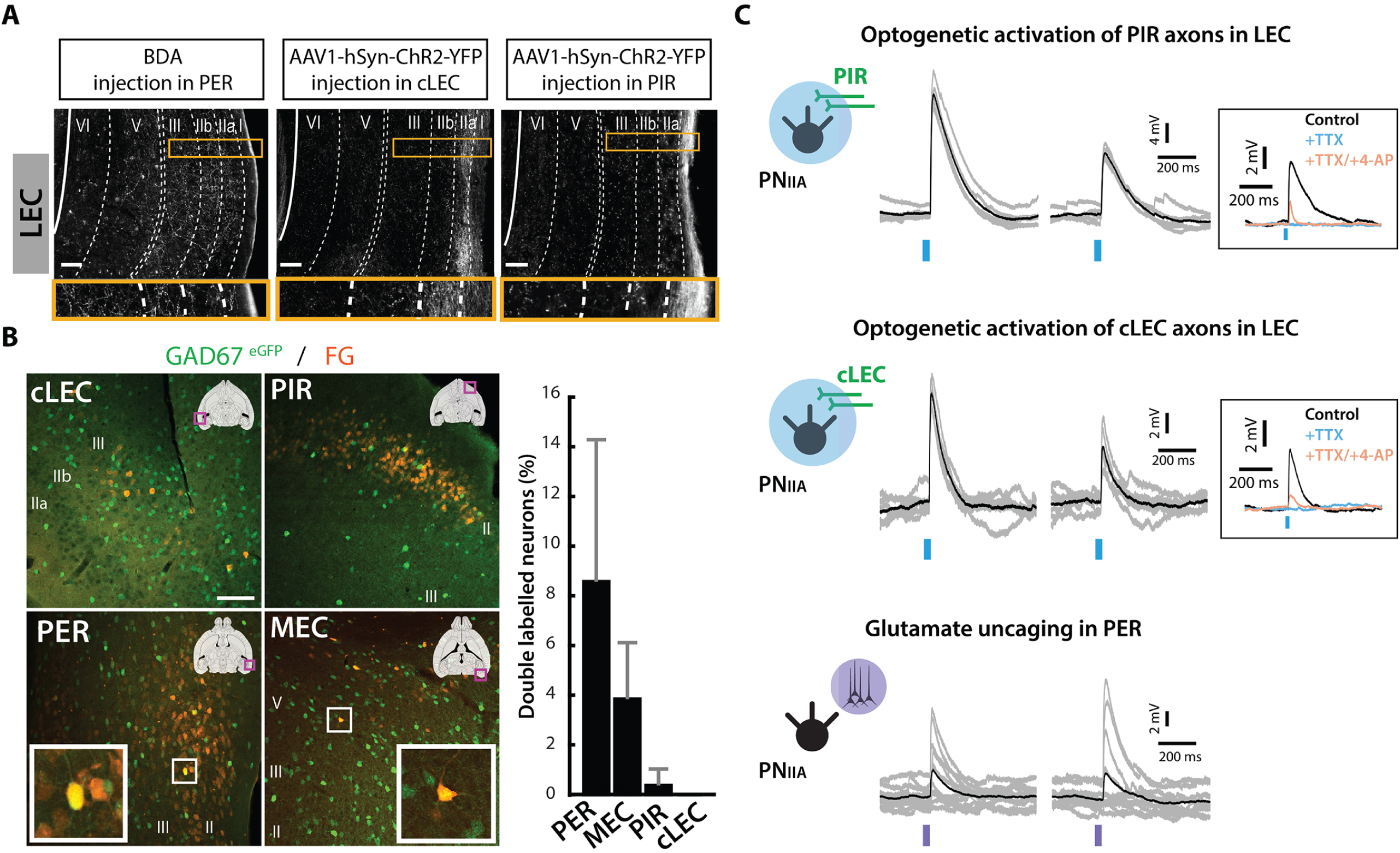
Cortical glutamatergic inputs to principal neurons in layer IIa. **(A)** Labelled axons in superficial layers of LEC following anterograde tracer injections into PER (left), cLEC (middle) and PIR (right). BDA, biotinylated dextran amine. See Supplementary Fig.11 for images of the injection sites. **(B)** Confocal images of neurons retrogradely labelled from LEC in a GAD67^eGFP^ mouse (same case as the experiment in **Fig. 1A**). FG^+^ cell bodies were found in MEC, PER, cLEC and PIR. In this brain, co-localization of GFP and FG was evident in PER and MEC (white square insets). Bar graphs show the proportions of neurons retrogradely labelled with FG or AAV2-CAG-tdTomato that also expressed GFP (FG/tdTomato and GFP; mean ± standard deviation, n = 3 mice). **(C)** Electrophysiological interrogation of afferent pathways in wild-type mice. Optogenetic activation (blue bars) of photosensitive axons from PIR (top) or cLEC (middle) in LEC elicited monosynaptic EPSPs in PN_IIA_. Insets show average membrane potential traces in control ACSF (black), and ACSF added with TTX (blue) or TTX + 4-AP (orange). Activation of PER neurons (bottom) through UV-photostimulation (purple bars) of caged glutamate evoked EPSPs in PN_IIA_ in LEC. All membrane potential traces show the average trace (black) superimposed on individual trials (grey). Note that the recordings in the three panels are from different mice. Scale bars are 100 µm in **(A)** and **(B)**.

To assess synaptic connectivity with PN_IIa_, in separate wild-type mice, we virally expressed ChR2 in axons from PIR or cLEC. Because of the proximity between PER and the recording location in dorsal LEC, we decided to drive PER activity through patterned laser scanning photostimulation of caged glutamate. Activation of these three pathways while *in vitro* current-clamp recording PN_IIa_ at their resting membrane potential (−67.8 [−69.8 – −66.0] mV; median [I QR]) elicited depolarizing potentials in responding neurons (PER, n = 8 neurons from 4 mice; PIR, n = 32 neurons from 8 mice; cLEC, n = 14 neurons from 6 mice; Fig. 4C). These depolarizing responses were classified as excitatory postsynaptic potentials (EPSPs) as they were observed at voltages positive to the calculated chloride equilibrium potential (≈-69 mV for our solutions). Although delayed inhibitory responses were occasionally observed following the initial EPSP, we did not detect exclusive laser-evoked hyperpolarizing responses. I t should be noted, however, that it is possible that inhibitory effects could have been masked in some recordings, due to the resting membrane potential in our recordings being close to the theoretical chloride equilibrium potential. This could be particularly relevant for the inputs from PER, an area where we found LEC-projecting GABAergic neurons (Fig. 4B) and which is known to provide long-distance inhibition to LEC (Pinto et al., 2006; Apergis-Schoute et al., 2007). A pharmacological test (+TTX/+4-AP) uncovered that optogenetic activation of axons from PIR or cLEC gave rise to monosynaptic responses in PN_IIa_ (PIR, 6 cells from 1 mouse; cLEC, 3 cells from 1 mouse; Fig. 4C, insets). Taken together, our electrophysiological and anatomical findings indicate that PIR, cLEC and PER provide a prevailing excitatory input to PN_IIa_.

Our observation that the population of PN_IIa_ receive synaptic inputs from PER, PIR, cLEC as well as MEC suggests that individual LEC PN_IIa_ may integrate synaptic input from several cortical areas. We therefore sought to test in the same experiment whether single PN_IIa_ responds to activation of two upstream brain areas. Synaptic convergence was tested by monitoring the membrane potential of recorded PN_IIa_ while, in sequence, activating neurons in PER and optogenetically activating axons from either PIR, cLEC or MEC (PER– PIR/ PER–cLEC/ PER–MEC). To increase the stimulation efficacy of neurons in PER, we replaced glutamate uncaging with extracellular electrical stimulation of the superficial layers of PER, in agreement with the experimental design of a previous study in the rat (de Villers-Sidani et al., 2004). We found that single PN_IIa_ that responded to PER stimulation often also responded to stimulation of axons from the other tested brain region (PER–PIR: 67.7%, n = 21/31 neurons; PER-cLEC: 77.3%, n = 17/22 neurons; PER–MEC: 59.5%, n = 25/42 neurons), confirming that perirhinal input converges with synaptic inputs from cLEC, PIR and MEC at the level of single PN_IIa_ in LEC (Supplementary Fig.12).

### Synaptic inputs from MEC diminish glutamatergic synaptic events in principal neurons in layer IIa

In a final set of experiments, we tested in 4 wild-type mice how inputs from MEC may modulate postsynaptic excitatory responses in PN_IIa_. To this end, we used a dual-color photostimulation approach in order to combine glutamate uncaging with virally-mediated (AAV1-hSyn-ChrimsonR-tdTomato) optogenetics. EPSPs were generated by focal UV-photolysis of caged glutamate at dendrites of PN_IIa_ while synaptic release from MEC was triggered by photostimulation of medial entorhinal ChrimsonR^+^ axons in layer I of LEC (Fig. 5A). This procedure allowed to examine whether synaptic potentials produced by MEC will add or subtract to the EPSPs elicited by photorelease of glutamate. Sequential photorelease of caged glutamate transiently depolarized PN_IIa_ (Fig. 5B), thereby mimicking EPSPs generated by afferent excitatory pathways targeting PN_IIa_ (e.g. axons from PIR, PER or cLEC). Pairing release of glutamate with photoactivation of MEC axons led to evoked EPSPs that were smaller (middle trace, Fig. 5B) than those generated before and after by photolysis of caged glutamate alone (left and right traces in Fig. 5B; Figs. 5C and 5D). Across the entire set of recordings, synaptic input from MEC reduced the average amplitude as well as the average duration of uncaging-evoked EPSPs (n = 38 neurons; amplitude: p< 0.0001; duration: p = 0.0078; Wilcoxon signed rank test; Figs. 5E–5G). Only in a single PN_IIa_ did we observe that postsynaptic potentials evoked by MEC and experimentally released glutamate summed to a larger amplitude EPSP. These data are consistent with our earlier observations of a strong MEC inhibitory input to PN_IIa_ (Figs. 2 and 3). Our observations further demonstrate that the MEC–LEC pathway can reduce the amplitude and duration of concomitant EPSPs arising from excitatory inputs targeting PN_IIa_ dendrites in layer I, suggesting that MEC may impose a shunting inhibitory effect on EPSPs produced by other afferent excitatory pathways targeting LEC PN_IIa_, such as those originating in PER, cLEC and PIR (Fig. 4).

**Figure 5.**
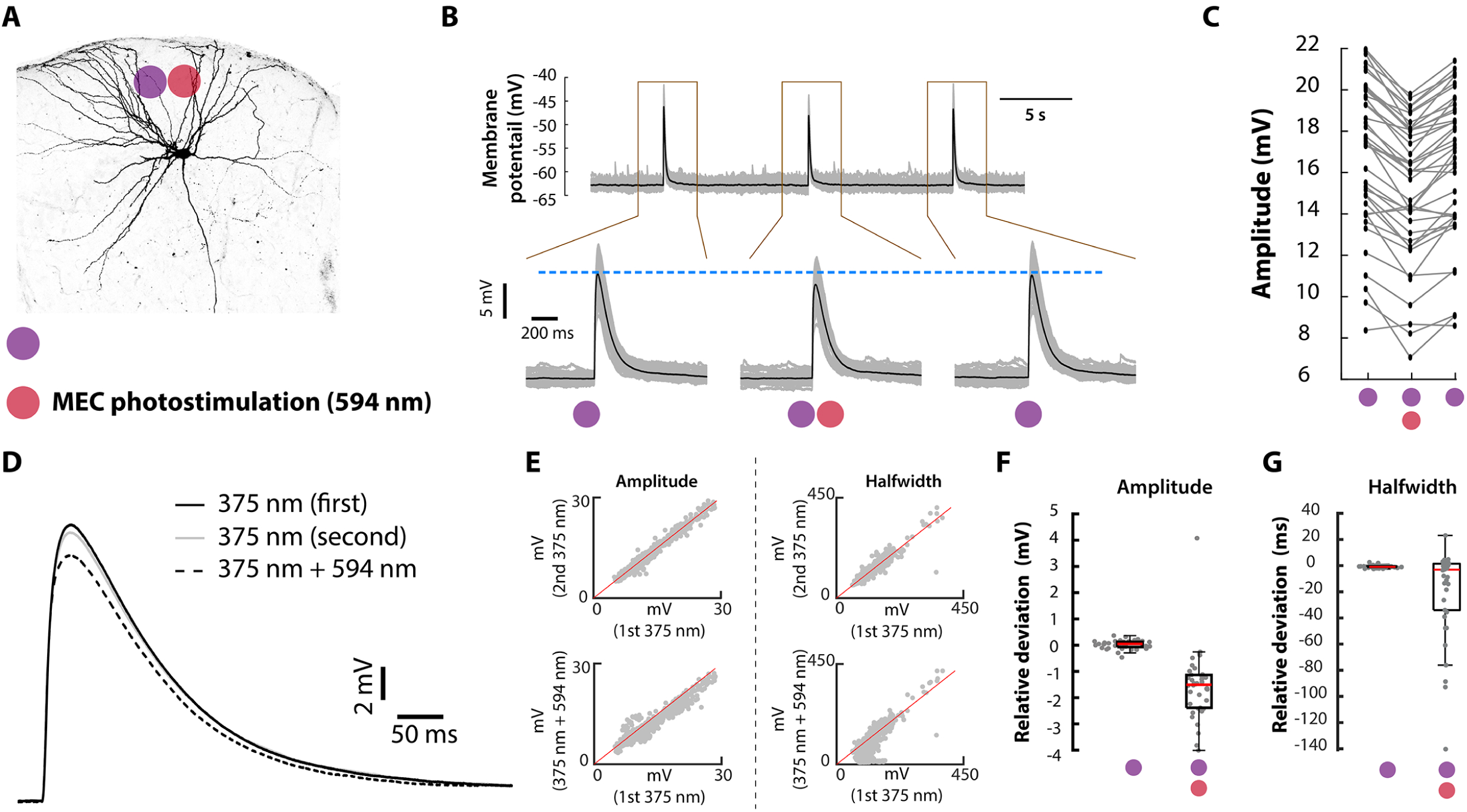
Synaptic inputs from MEC diminish EPSPs of principal neurons in layer IIa. **(A)** Experimental design of dual photostimulation within layer I of LEC while recording from PN_IIA_. Depicted is an example of a biocytin-filled PN_IIA_ showing its typical dendritic arborizations in layers I/IIa. **(B)** Example PN_IIA_ recording showing EPSPs evoked by glutamate uncaging alone (left, right) or together with MEC input (middle). The average membrane potential trace (black) is superimposed on the individual traces (grey). The dotted blue line indicates the peak of the average EPSP in response to photolysis of glutamate alone, showing a decrease in EPSP amplitude when pairing glutamate release with MEC activation. **(C)** Quantification of EPSP amplitude for all individual trials recorded from the neuron in **B**. **(D)** Average voltage traces from the recording in **B**. **(E)** Scatter plots of EPSP amplitude (left) and halfwidth (right) for individual sweeps of the experimental protocol from all recorded PN_IIA_ (n = 38). **(F)** Relative deviations in EPSP amplitude between responses elicited when glutamate was released alone (purple) and when glutamate release was paired with photostimulation of MEC axons (purple and red). Data show relative deviations for all recorded PN_IIA_ (n = 38). **(G)** Same as in **F**, but for EPSP halfwidth.

## DISCUSSION

We have used a combination of viral tracing and optogenetic-assisted circuit mapping to identify a functional circuit connecting MEC to neurons located in the superficial layers of LEC, which supply the main lateral entorhinal input to the hippocampal formation. A main finding of our work is that MEC contains a population of SST^+^ GABAergic neurons that provide dense and restricted axonal innervation of the most superficial layers (layer I and IIa) of LEC. Optogenetic stimulation of these axons gave rise to strong inhibition, as revealed by the synaptic charge transfer, among recorded principal cells in layer IIa. This contrasted with weaker responses among principal neurons in layers IIb and I II. This result was consistent between optogenetic experiments where inhibitory synaptic inputs were pharmacologically isolated and optogenetic experiments in slices from transgenic SST^cre^ mice where only axons from MEC SST^+^ neurons were activated. A strong targeting of PN_IIa_, which mainly constitute morphologically defined fan cells, by MEC SST^+^ axons is supported by anatomical data showing that the dendritic tree of fan cells almost exclusively distribute in layers IIa and I (Tahvildari and Alonso, 2005; Canto and Witter, 2012; Leitner et al., 2016; Nilssen et al., 2018; Vandrey et al., 2022), and is thus well-positioned to overlap with afferent MEC SST axons. Although PN_IIb_ and PN_III_ have apical dendrites in layers IIa/I, a possible severing of dendrites during brain slice preparation is unlikely to explain the weaker responses observed among these neurons, because *post hoc* visualization of recorded PN_IIb_/PN_III_ revealed many intact dendrites in layers IIa/I. Taken together with our data showing that local GABAergic neurons in LEC receive only sparse inhibitory input from MEC SST neurons, the findings suggest that PN_IIa_ is the preferred postsynaptic partner in superficial LEC of MEC long-range SST^+^ axons. It should be noted that principal neurons in the deeper layers, V and VI, whose apical dendrites also extend into layer I, are additional possible targets of MEC SST axons (Canto and Witter, 2012; Ohara et al., 2021), however, this was not tested in the present study.

Even though long-range projecting GABAergic axons are found in several pathways linking brain areas supporting spatial navigation and memory (Melzer and Monyer, 2020), the pathways connecting the two divisions of the EC have so far been considered to be purely excitatory. In contrast to the clear GABAergic innervation of LEC by MEC SST neurons, we did not find evidence for an SST^+^ GABAergic component in the pathway connecting LEC to MEC. This suggests that long-range inhibition from LEC to MEC is either absent or mediated by inhibitory neuron types other than SST^+^ neurons. Long-range projecting GABAergic neurons are present in LEC and have been found to innervate the hippocampus (Basu et al., 2016; Leitner et al., 2016); however, the molecular identity of these neurons is not known. Additional tracing experiments are needed to map out a possible innervation of MEC by LEC long-range GABAergic neurons belonging to those expressing either the molecular markers parvalbumin or the type 3A serotonin receptor, which together with SST neurons make up the majority of LEC GABAergic neurons (Leitner et al., 2016).

The observation that LEC PN_IIa_ are main targets of MEC inhibitory axons points to a circuit wiring that is different from most other inhibitory pathways so far described in the greater hippocampal network. With the exception of the projection from the HF to the retrosplenial cortex (Yamawaki et al., 2019), long-range inhibitory axons preferentially synapse onto local inhibitory neurons within the target area, as shown for inhibitory pathways connecting MEC with HF in both directions (Melzer et al., 2012), the medial septum to MEC (Gonzalez-Sulser et al., 2014; Unal et al., 2015; Fuchs et al., 2016), to LEC (Desikan et al., 2018) and HF (Freund and Antal, 1988), as well as the presubiculum to MEC (van Haeften et al., 1997). Additionally, long-range axons from GABAergic neurons in LEC have been shown to innervate hippocampal interneurons (Basu et al., 2016). Activation of such circuits supplying direct GABAergic input to interneurons has been proposed to have an excitatory effect on principal neurons by releasing them from inhibition (Melzer and Monyer, 2020). In contrast, the present data demonstrate a circuit producing clear monosynaptic inhibition of excitatory PN_IIa_ in LEC, suggesting a predominant inhibitory influence. Although further study is required to establish the exact functional relevance in vivo of the inhibitory MEC to LEC circuity, our data point to a possible role of MEC in suppressing output from LEC to the hippocampus by directly inhibiting the activity of PN_IIa_. Furthermore, we showed that the MEC–LEC pathway can reduce the amplitude and duration of concomitant EPSPs arising from excitatory inputs targeting PN_IIA_ dendrites in layer I. Our data, taken together with previously published data indicate that layer I excitatory inputs predominantly originate from intrinsic ipsilateral and contralateral projecting neurons in LEC (Leitner et al., 2016; Ohara et al., 2019), as well as from piriform and perirhinal cortex. This implies that integration over time of these inputs, as well as their efficacy to induce action potentials in PN_IIA_ will be reduced,

In addition to long-distance SST^+^ axons, our data show that MEC sends a dense projection of glutamatergic excitatory axons to LEC. Consistent with earlier anatomical descriptions of MEC innervation of LEC (Köhler, 1986; Witter et al., 1986; Dolorfo and Amaral, 1998; Chrobak and Amaral, 2007; Ohara et al., 2019), these axons distribute across several layers of LEC and derive from neurons in several layers of MEC. Our data from both voltage clamp and current clamp recordings indicate that MEC axons provide stronger excitatory input to neurons in layers IIb and I II than layer I Ia neurons. This pattern differs from the connectivity from olfactory areas to LEC. Consistent with the present data showing strong excitatory inputs to LEC PN_IIa_ following optogenetic activation of PIR axons, another recent study found that axons from PIR as well as the olfactory bulb target principal neurons in layer IIa more strongly than principal neurons in layer IIb (Bitzenhofer et al., 2022). I t remains for future experiments to identify the neuron populations in MEC that provide excitatory input to principal neurons in superficial layers of LEC. I t is likely that the excitatory responses observed in this study, at least in part, reflect input from molecularly defined calbindin-expressing principal neurons in MEC layer II, which have been shown to innervate superficial layers of LEC (Ohara et al., 2019). In contrast to LEC PN_IIa_ which send axons to MEC in addition to the hippocampus (Vandrey et al., 2022), hippocampal-projecting stellate neurons in MEC layer II do not project to LEC (Nilssen et al., 2018), which highlights a salient difference in the organization of the intra-entorhinal pathways (MEC to LEC and LEC to MEC).

A traditional view of the functional organization of the hippocampal-parahippocampal system suggests that distinct information is represented by LEC and MEC and relayed by largely segregated processing streams to the hippocampus, where the information is further processed and integrated (e.g. Eichenbaum et al. 2012). Together with previous studies showing that MEC and LEC are interconnected (Köhler, 1986; Witter et al., 1986; Dolorfo and Amaral, 1998; Biella et al., 2002; Chrobak and Amaral, 2007; Ohara et al., 2019; Vandrey et al., 2022), the present data speak against clearly separate processing streams through LEC and MEC and instead suggest circuits through which information represented in LEC and MEC could be integrated already at the level of the EC. In particular, the present study together with that of Vandrey et al. (2022) emphasize that MEC and LEC are connected in ways that allow entorhinal output to the hippocampus, originating from one subdivision, to be directly influenced by the other entorhinal subdivision. In our data, we noticed that MEC GABAergic and glutamatergic axons target postsynaptic neurons in a manner that matches the different connectivity patterns of LEC with the HF (Figure 6). MEC SST^+^ axons readily connect with PN_IIa_, which provide input to DG-CA2/CA3 (Steward and Scoville, 1976; van Groen et al., 2003; Kohara et al., 2014; Leitner et al., 2016; Lopez-Rojas et al., 2022), whereas MEC glutamatergic axons preferentially synapse onto neurons in layers IIb and III, which issue projections to CA1 and subiculum (van Groen et al., 2003; Kitamura et al., 2014; Masurkar et al., 2017; Ohara et al., 2019). Although the HF is generally considered as a main target of axons of PN_IIa_ and PN_III_ (Witter et al., 2017), the synaptic targets of neurons in layer IIb are reportedly more diverse and include MEC and olfactory areas, in addition to CA1 (Leitner et al., 2016; Ohara et al., 2019).

**Figure 6.**
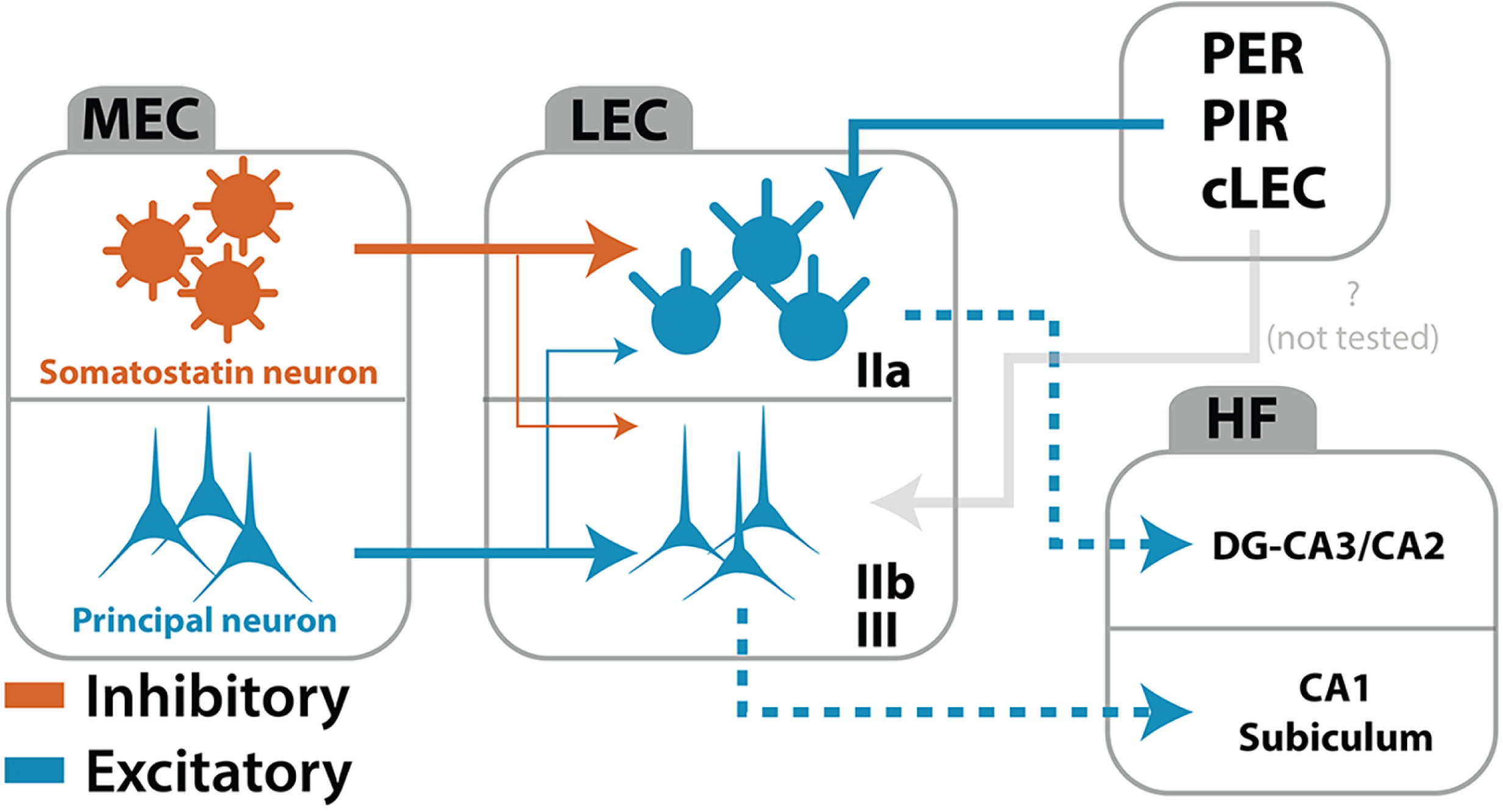
Anatomical organization of the circuit connecting MEC to LEC. Inhibitory and excitatory pathways differentially target principal neurons in the superficial layers of LEC. The main pathways (solid lines) described in this study are shown together with the well-established connectivity between LEC and the hippocampus (dotted lines; Nilssen et al., 2019). For the MEC–LEC pathway, the thickness of the GABAergic/glutamatergic arrows indicate the relative strength of these inputs onto postsynaptic neurons. HC, hippocampus

The organization of MEC synaptic inputs to LEC as described in this study has two main implications for how MEC might modulate lateral entorhinal communication with the HF (Figure 6). First, the weaker relationship between MEC glutamatergic axons and PN_IIa_ as compared to PN_IIb_ and PN_II_ indicates that activity in the glutamatergic pathway would impose a weaker excitatory influence on lateral entorhinal neurons projecting to DG-CA3/CA2 than on neurons projecting to CA1 and subiculum. Second, the preferred synaptic inhibition onto PN_IIa_ mediated by MEC SST neurons suggests that lateral entorhinal output to DG and CA2-CA3, rather than output to CA1 and subiculum, is under powerful and direct MEC-derived inhibitory influence. The organization of synaptic inputs to LEC neurons is consistent with the possibility that MEC may route output from LEC by modulating the efficacy by which signals travel to downstream regions of the HF, either through a suppression of activity in the pathway from layer IIa to DG-CA3/CA2 or a facilitation of activity in the pathway from layers IIb/III to CA1 and subiculum. The relevance of this organization is yet to be determined, both with respect to effects on intrinsic hippocampal circuits, as well as taking into account data indicating that inputs to PN_IIA_, and PN_III_ may differ in that inputs from medial prefrontal cortex, amygdala, and insular cortex preferentially seem to target PN_III_ neurons (Ohara et al., 2025; Ohara et al., 2014; Aléman-Andrade et al., 2025).

## MATERIALS AND METHODS

### Animals

We used transgenic B6.129P2-Pvalb*tm*1(*cre*)*Arbr*/J (PV^Cre^; RRID:IMSR_JAX:017320, The Jackson Laboratory), Sst*tm*2.1(*cre*)*Zjh*/J (SST^Cre^; RRID:IMSR_JAX:013044, The Jackson Laboratory) and Vip*tm*1(*cre*)*Zjh*/J (VIP^Cre^; RRID:IMSR_JAX:010908, The Jackson Laboratory) mice, GAD67*^eCFP^* mice (bred in-house; Tamamaki et al. 2003) and wild-type mice (C57BL/6N, RRID:IMSR_CRL:027, Charles River Germany). Both female and male mice were used and randomly selected. All mice were older than 6 weeks at the time of the experiment. Animals were maintained on a 12 hour light / 12 hour dark schedule, and group-housed in transparent acrylic cages with ad libitum access to food and water. All experiments were performed in accordance with the Norwegian Animal Welfare Act and the European Convention for the Protection of Vertebrate Animals used for Experimental and Other Scientific Purposes. The experiments were approved by the Norwegian Food Safety Authority (permit no. 8083 and 22312).

### Surgeries

Anesthesia was induced by placing the mouse in a chamber filled with isoflurane (5%, 1.2 L/min flow rate; Nycomed Zurich), after which it was mounted in a stereotaxic frame (Kopf Instruments). The animal was placed on a heating pad (37 °C) to maintain stable body temperature and protective ointment was applied to the eyes. Before the start of surgery, the mouse was given subcutaneous injections of buprenorphine hydrochloride (Temgesic®, 0.1 mg/kg; Indivior) and meloxicam (1 mg/kg; Metacam®, Boehringer Ingelheim Vetmedica), in addition to bupivacaine hydrochloride (1 mg/kg; Marcain™, Astra Zeneca) at the surgical incision site. Anterior-posterior coordinates were measured from bregma (APb) or the posterior transverse sinus (APt), mediolateral (ML) coordinates were measured from the mid-sagittal sinus, and dorso-ventral (DV) coordinates were measured from the dura. Injection sites for all four areas (MEC, cLEC, PIR and PER) were derived from a stereotaxic mouse brain atlas (Franklin and Paxinos, 2007) and adjusted based on an extensive in-house collection of tracing experiments. The following coordinates (in mm) were used: LEC: +1.9 (APt), +4.75 (ML), −2.2 (DV); MEC: +0.50 (APt), +3.10 (ML), −1.8 (DV); PIR: −0.6 (APb), +4.4 (ML), −3.1 (DV); PER: +2.1 (APt), +4.6 (ML), −1.2 (DV). After stereotactic injections, the wound was cleaned and sutured, and the animal was put back in its home cage. All injections were performed using glass micropipettes. Pressure injections used an automated microinjection pump (Nanoliter 2010, World Precision Instruments) connected to a microsyringe pump controller (Micro4 pump, World Precision Instruments). Iontophoretic injections used a current source driving alternating 6 seconds on/off current (5-15 minutes; 7 µA or 6 µA; Stoelting Midgard).

### Anterograde and retrograde chemical tracer injections

Wild-type mice received a single iontophoretic injection of each anterograde tracer: biotinylated dextran amine (BDA; 10 kDa; 5% solution in 0.125 M phosphate buffer, pH 7.4; Thermo Fisher Scientific, Cat. No. D1956) or Phaseolus vulgaris leucoagglutinin (PHA-L; 2.5% solution in 10 mM phosphate buffer, pH 7.4; Vector Laboratories, Cat. No. L-1110-5). To reduce the number of animals, the two tracers were deposited in two different cortical areas, i.e., MEC, PER, cLEC or PIR, in the same animal. The retrograde neuroanatomical tracer fluorogold (FG; 2.5% in H2O, Fluorochrome, ID: Fluorogold) was pressure injected into dorsal LEC, directly ventral to the rhinal fissure, of GAD67*^eCFP^* mice.

### Virus injections

Each virus was deposited via pressure injection. All virus injections were unilateral. In all cases, 100-300 nL of the virus solution was delivered at 40 nL/min.

#### Wild-type mice

For optogenetic experiments or anterograde viral tracing in wild-type mice, an injection of AAV1 carrying a fused channelrhodopsin2-YFP protein (AAV1-hSyn-ChR2-H134R-eYFP-WPRE, Addgene, viral prep no. 26973-AAV1) was placed into either MEC, LEC or PIR. For dual-color optogenetic and uncaging experiments, mice were injected with AAV1-hSyn-ChrimsonR-tdTomato (Addgene, viral prep no. 59171-AAV1) into MEC.

#### SST^Cre^ mice

In SST^Cre^ mice used for optogenetic experiments, AAV1-hSyn-FLEX-ChrimsonR-tdTomato (UNC vector core, deposited by Dr. Ed Boyden) was injected into MEC. Virus-mediated anterograde tracing from MEC used injections of either AAV1-CAG-FLEX-tdTomato (Addgene, viral prep no. 28306-AAV1) or AAV9-CAG-FLEX-GFP (UNC vector core, deposited by Dr. Ed Boyden), or a mixture consisting of equal amounts of AAV1-CAG-FLEX-tdTomato and AAV1-CamKII -eGFP-WPRE-rBG (Addgene, viral prep no. 105541-AAV1). To specifically label SST neurons in LEC, mice received a single injection of AAV1-CAG-FLEX-tdTomato into LEC. For monosynaptic rabies virus tracing, mice were first injected with a helper virus (AAV5-hSyn-FLEX-splitTVA-2A-mCherry-2A-B19G; produced in-house, modified after Addgene plasmid no. 52473) into MEC, and after two weeks they received an injection of an in-house manufactured EnvA pseudotyped G-protein deleted rabies virus expressing GFP in the same anatomical location (see Jacobsen 2019 for details regarding virus production).

#### PV^Cre^ and VIP^Cre^ mice

To selectively label MEC VIP^+^ or PV^+^ neurons, AAV1-CAG-FLEX-tdTomato or AAV5-hSyn-FLEX-splitTVA-2A-mCherry-2A-B19G was injected into MEC of VIP^Cre^ or PV^Cre^ mice, respectively.

#### GAD67^eGFP^ mice

For optogenetic interrogation of MEC inputs to LEC, AAV1-hSyn-ChrimsonR-tdTomato was injected into MEC. To retrogradely label neurons projecting to LEC, AAV2-CAG-tdTomato (Addgene, viral prep no. 59462-AAV2) was injected into LEC.

### Preparation of acute brain slices

Mice of either sex (6-16 weeks old) were used for electrophysiological experiments. For virus-injected animals, acute brain slices were collected 2-4 weeks after surgery. Mice were kept under isoflurane anesthesia until reflexes were no longer observed, after which they were killed by decapitation. The brain was extracted from the skull and submerged in ice-cold (≈0 °C), continuously oxygenated (95% O_2_/5% CO_2_), cutting artificial cerebrospinal fluid (ACSF). The cutting ACSF had a composition of 110 mM choline chloride, 2.5 mM KCl, 25 mM D – Glucose, 25 mM NaHCO_3_, 11.5 mM sodium ascorbate, 3 mM sodium pyruvate, 1.25 mM NaH_2_PO_4_, 100 mM D – Mannitol, 7 mM MgCl_2_ and 0.5 mM CaCl_2_. Osmolality and pH were adjusted to 430 mOsm and 7.4, respectively. The brain hemisphere ipsilateral (for viral injections into PIR or MEC) or contralateral (for viral injections into LEC) to the virus injection site was glued onto the stage of a vibratome (Leica VT1000S, Leica Biosystems). Semicoronal slices (350 µm thickness, 20° tilt with respect to the coronal plane) were acquired. This cutting angle was chosen to preserve the dendritic tree of principal neurons in superficial layers of LEC (Tahvildari and Alonso, 2005; Canto and Witter, 2012) as well as to secure axonal projections from PER to LEC (de Villers-Sidani et al., 2004). Slices were immediately transferred to a heated chamber (35 °C) and submerged in oxygenated holding ACSF (high Mg^2+^ – low Ca^2+^) solution containing 126 mM NaCl, 3 mM KCl, 1.2 mM Na_2_HPO_4_, 10 mM D-glucose, 26 mM NaHCO_3_, 3 mM MgCl_2_ and 0.5 mM CaCl_2_. Slices were maintained at 35°C for 30 minutes and then allowed to adjust to room temperature until use.

### Whole-cell patch-clamp recordings

The slices were moved to a recording rig containing a Axio Examiner.D1 microscope (Carl Zeiss) equipped with a 20x / 1.0 NA water immersion objective and infrared differential interference contrast optics. Patch pipettes (resistance: 2-8 MΩ) were prepared from borosilicate glass capillaries (1.5 outer diameter x 0.86 inner diameter; Harvard Apparatus) using a Flaming/Brown micropipette puller (P-97; Sutter Instruments) and filled with the appropriate intracellular solution (see below). Patch clamp recordings were performed at 35°C and slices were superfused with oxygenated recording ACSF containing 126 mM NaCl, 3 mM KCl, 1.2 mM Na_2_HPO_4_, 10 mM D-Glucose, 26 mM NaHCO_3_, 1.5 mM MgCl_2_ and 1.6 mM CaCl_2_. Somata of up to four neurons were targeted for simultaneous recordings.

Following the formation of a gigaohm resistance seal between the pipette and the cell membrane, pipette capacitance compensation was performed, and the cell membrane was subsequently ruptured to obtain whole-cell recording configuration. Data were acquired with an EPC 10 Quadro USB amplifier (Heka Elektronik), with amplifier commands and data acquisition controlled by Patchmaster (version 2.90, Heka Elektronik). Acquired data were low-pass (bessel) filtered at 4 kHz and digitized at 10 kHz. In a subset of experiments, bicuculline (10 µM, Sigma-Aldrich, Cat. No. 14343) was introduced to the circulating ACSF to block GABA_A_-mediated synaptic transmission, whereas combined application of DNQX (10 µM, Tocris Bioscience, Cat. No. 0189) and DL–APV (50 µM, Tocris Bioscience, Cat. No. 0105) was used to block glutamatergic synaptic transmission. Sequential addition of tetrodotoxin (TTX, 1 µM, Tocris Bioscience, Cat. No. 1078) followed by a mixture of both 4-Aminopyridine (4-AP, 100 µM, Sigma-Aldrich, Cat. No. 275875) and TTX was carried out to test whether laser-evoked events reflected monosynaptic inputs (Petreanu et al., 2009).

### Voltage clamp

Whole-cell voltage clamp recordings used a cesium-based internal pipette solution (117 mM cesium gluconate, 13 mM CsCl, 2 mM MgCl_2_, 10 mM HEPES, 10 mM Na_2_-phosphocreatine, 0.3 mM Na – GTP, 4 mM Mg – ATP, 5 mM QX314-Cl). Biocytin (I ris Biotech, Cat. No. LS-3510) was added at a concentration of 5–8 mg/mL. Series resistance was continually monitored and maximally compensated (i.e., kept slightly below the compensation level where oscillations start to occur). Recordings with series resistance ≤ 25 MΩ were accepted (16.2, 4.9 – 24.9 MΩ; mean, range). In all experiments, we recorded inhibitory postsynaptic currents (IPSCs) and excitatory postsynaptic currents (EPSCs) by clamping the membrane potential to the reversal potential for excitatory currents (E_exc_ ≈0 mV) and inhibitory, chloride-carried currents (E_Cl−_ ≈-50 mV), respectively. The holding potentials were corrected for the liquid junction potential (≈10 mV as measured experimentally).

### Current clamp

Whole-cell current clamp recordings used a potassium gluconate-based intracellular solution of the following composition: 120 mM K – gluconate, 10 mM KCL, 10 mM Na_2_-phosphocreatine, 10 mM HEPES, 4 mM Mg-ATP, 0.3 mM Na-GTP, with pH adjusted to 7.3 and osmolality to 300-305 mOsm. Biocytin (5–8 mg/mL) was added to the internal solution. All recordings (series resistance: 18.9 MΩ, 5.0 – 66.7 MΩ; mean, range) were included for analysis. Online bridge balance adjustments were performed. No correction was carried out for the liquid junction potential (≈13 mV as measured experimentally). In one set of experiments (Fig. 3), a depolarizing step current (500 ms duration) was injected through the pipette into the soma to induce firing of action potentials. The intensity of the applied current was adjusted to the weakest current required to reach threshold and elicit firing of 1-7 action potentials.

### Optogenetic activation of photosensitive axons

Laser-scanning photostimulation was accomplished using a UGA-42 GEO point scanning system (Rapp OptoElectronic) equipped with three continuous laser lines (375 nm, 473 nm and 594 nm), and controlled by SysCon software (v. 1.1.8.0, Rapp OptoElectronic). For optogenetic activation of ChR2 or ChrimsonR, the 473 nm or 594 nm diode laser line was used, respectively. The intensity of laser output from the objective was set to ≈8 mW in all experiments using optogenetic activation of ChR2 through injections of AAV1-hSyn-ChR2-H134R-eYFP-WPRE into MEC. To photoactivate virally-expressed ChrimsonR, the laser intensity was set to ≈3 mW (for GAD67*^eCFP^* experiments) or ≈4 mW (for SST^+^ experiments). In mice with injections of AAV1-hSyn-ChR2-H134R-eYFP-WPRE into PIR or cLEC, laser intensity (1.5–5.0 mW) was adjusted for each simultaneously recorded neuron cluster, depending on the virus expression levels in the slice, to evoke sub-threshold membrane potential deflections. Laser stimuli were confined to a single location (pulse duration: 100 ms [473 nm]; 10 ms [594 nm]), or they were arranged in a grid pattern consisting of several equidistant stimulation positions (473 nm: one row containing seven stimulation spots or four rows à five spots, pulse duration was 1 ms; 594 nm: one row containing seven stimulation spots, pulse duration was 5 ms). Stimuli were delivered at a frequency of 2 Hz. All 473 nm and 594 nm laser stimuli had a beam diameter of ≈50 μm (light scattering in the medium not taken into account).

### Photostimulation of caged glutamate

MNI-caged-L-glutamate (Tocris Bioscience, Cat. No. 1490) was dissolved in 20 mL recording ASCF to a final concentration of 200 µM and recirculated. The solution was replaced when control responses evoked by uncaging near the cell bodies of recorded neurons started to decline in amplitude, or after 3 hours of recirculation (Bendels et al., 2008). Laser-scanning photostimulation of caged glutamate was performed with the same UGA-42 equipment as described above, except here a UV laser (375 nm) was used as light source. The beam diameter of the 375 nm laser stimuli was ≈1 µm. For glutamate uncaging in PER, laser stimuli were administered in a grid pattern (10 x 12 or 21 x 21 stimulation positions), with duration of 5 ms and intensity set to ≈8-9 mW. For uncaging directly at dendrites of principal neurons in layer IIa of LEC, glutamate was released by three repeating laser uncaging events of identical duration (5 ms) and intensity (10 seconds interstimulus interval), with stimulation intensity adjusted for each neuron in order to evoke subthreshold EPSPs. In these experiments, a single pulse of 594 nm laser (50 µm diameter, 10 ms duration, ≈4 mW intensity) was given 1 ms after the end of the second 375 nm illumination in order to photoactivate ChrimsonR-expressing MEC axons.

### Electrical stimulation of perirhinal cortex

Extracellular stimulation of perirhinal cortex was performed with a tungsten bipolar electrode (tip separation: 150-300 μm). The extracellular stimulation electrode was in all cases positioned in the superficial layers of PER adjoining LEC layer IIa, in accordance with anatomical data describing the main projection to dorsolateral LEC originating from layers II and III of PER (Burwell and Amaral, 1998). A single pulse stimulation (100 μs, 0.3-3 mA) was generated by an I so-Flex isolation unit (AMPI, Israel) controlled by Patchmaster. The stimulation intensity was adjusted for each recording to the minimum current required to evoke postsynaptic potentials in at least one of the simultaneously recorded cells. Experiments in which electrical stimulation failed to produce responses in any of the simultaneously recorded neurons indicated unsuccessful preservation of PER to LEC connectivity, and these were consequently excluded from analysis. The stimulation protocol was repeated 20-50 times.

### Immunohistochemistry and imaging

#### Slices from patch-clamp experiments

Slices were after patch-clamp recordings exposed to 4% paraformaldehyde (PFA, pH 7.4, Sigma-Aldrich, Cat. No. 16005) for 48 hours at 4 °C, and then rinsed in phosphate buffer (PB, 0.125 M, 4 × 10 min) at room temperature followed by permeabilization in PB containing 0.5% Triton X-100 (PB-Tx, 3 x 15 min). Next, slices were pre-incubated at room temperature for 2 hours in a blocking medium composed of PB-Tx and Normal Goat Serum (NGS, 5% or 10%, Abcam, Cat. No. ab7481), after which they were incubated with primary antibodies (72 hours at 4 °C; Table 1) in PB-Tx solution.

**Table 1.** Primary antibodies used in the study.

| **Antibody** | **Conc.** | **Supplier** | **Cat. No** | **RRID** |
| --- | --- | --- | --- | --- |
| Chicken anti-GFP | 1:500 | Abcam | ab13970 | AB_300798 |
| Rabbit anti-2A | 1:200 | Sigma-Aldrich | ABS31 | AB_11214282 |
| Rabbit anti-calbindin | 1:3000 | Swant | CB38 | AB_10000340 |
| Rat anti-SST | 1:200 | Sigma-Aldrich | MAB354 | AB_2255365 |
| Rat anti-RFP | 1:500 | ChromoTek | 5F8 | AB_2336064 |
| Guinea pig anti-NeuN | 1:1000 | Sigma-Aldrich | ABN90P | AB_2341095 |
| Mouse anti-reelin | 1:1000 | Sigma-Aldrich | MAB5364 | AB_11212203 |
| Mouse anti-mCherry | 1:500 | Clontech | 632543 | AB_2307319 |
| Goat anti-PHA-L | 1:1000 | Vector Laboratories | AS-2224-1 | AB_2315136 |

The sections were washed (PB-Tx, 4 x 15 min) and incubated for 24 hours at room temperature in a Pb-Tx solution containing the relevant secondary antibodies (Table 2). Depending on the kind of virally- or genetically encoded fluorescent tag already present in the sample, enhancement of GFP (for GFP/YFP samples) or red fluorescent protein (RFP; for tdTomato samples) was performed. To reveal biocytin-filled neurons, streptavidin conjugated to an Alexa Fluor™ (AF) fluorescent tag (1:400; Thermo Fisher Scientific; AF405 [Cat. No. S32351], AF546 [Cat. No. S11225] or AF633 [Cat. No. S21375]) was added together with the secondary antibodies. In a few samples, NeuroTrace™ 640/660 deep-red fluorescent Nissl (1:200; Thermo Fisher Scientific, Cat. No. N21483) accompanied the secondary antibodies as a cytoarchitectural marker instead of NeuN.

**Table 2.**
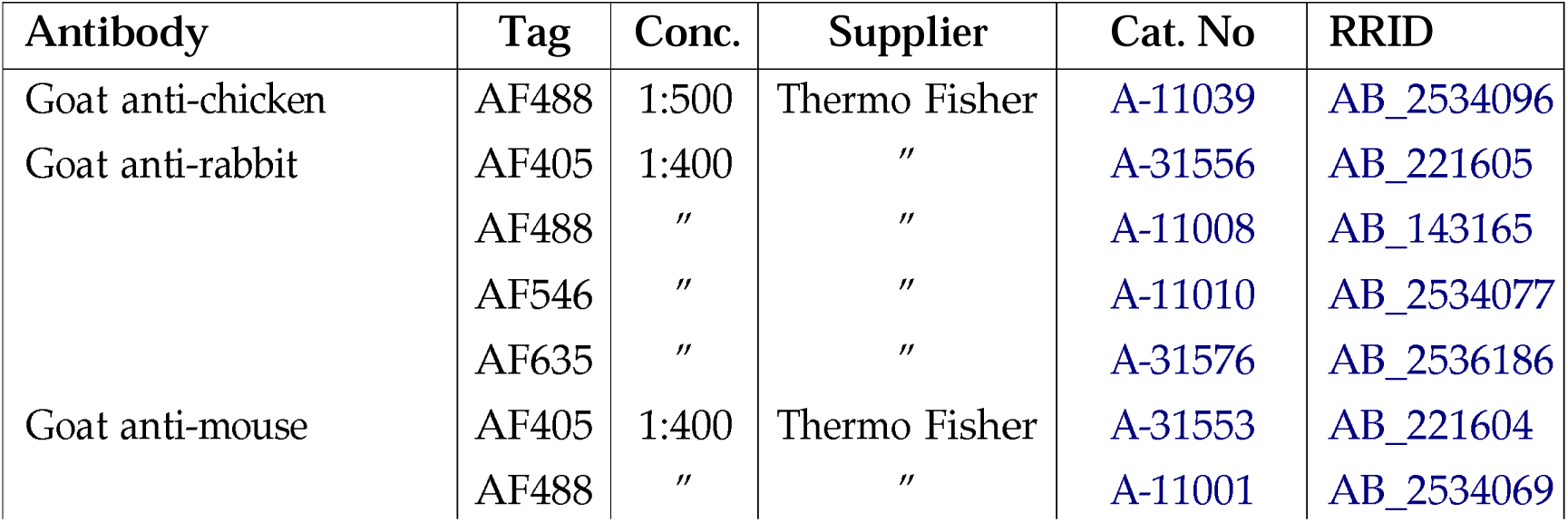
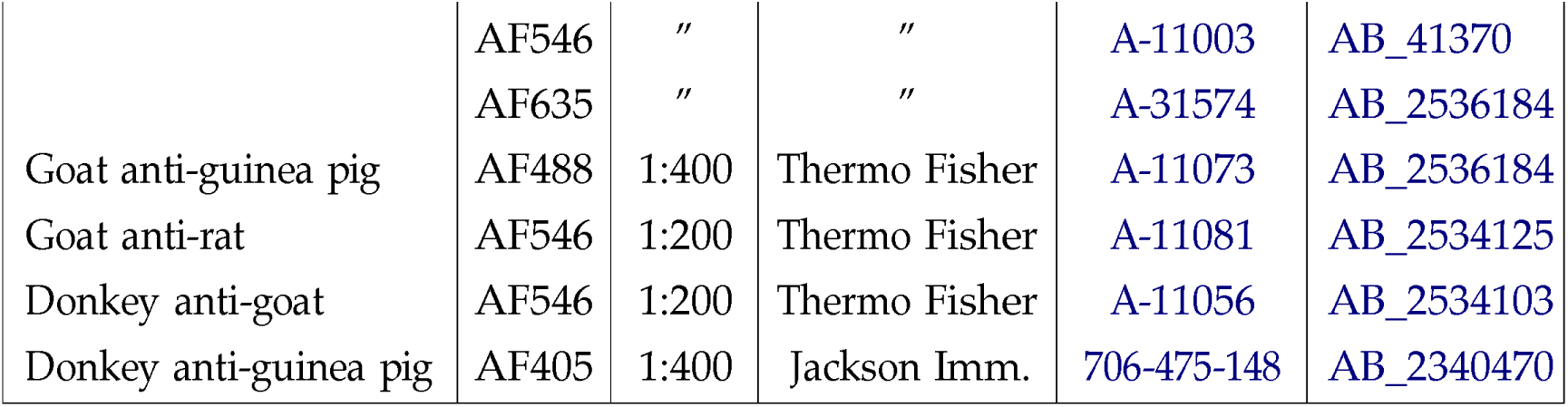
Secondary antibodies used in the study.

In preparation for confocal microscopy, the slices were first washed in PB (4 × 10 min) at room temperature and dehydrated through a series of increasing ethanol concentrations (30%, 50%, 70%, 90%, 100%, 100%, 10 minutes each). The slices were immersed in a mixture of 100% ethanol and methyl salicylate (1:1, 10 minutes) before being stored in methyl salicylate (VWR Chemicals). Some slices were stored in a cryoprotective solution containing glycerol (20%; VWR Chemicals) and dimethyl sulfoxide (2%; DMSO; VWR Chemicals) without prior dehydration. Slices were mounted in custom made metal slides with the storage solution and coverslipped before being imaged with a laser scanning confocal microscope (LSM 880 AxioImager Z2, Carl Zeiss). Image z-stacks were acquired with Plan-Apochromat objectives (10x, NA 0.45; 20x, NA 0.8) to retrieve the full morphology and location of the biocytin-filled recorded neurons, along with other immunomarkers present in the slice (NeuN/Neurotrace, GFP/tdTomato and reelin/calbindin). AF405 was excited with a 405 diode laser (bandpass emission filter: 410 – 461 nm), AF488 and GFP were excited by the 488 line of an Argon laser (bandpass emission filter: 490 – 543 nm), AF546 and tdTomato/RFP by a DPSS 568 laser line (bandpass emission filter: 570–623 nm), and AF633 by a HeNe 633 laser line (bandpass emission filter: 635-735 nm).

#### Slices from neuroanatomical tracing experiments

After 7–14 days of survival, virus- or tracer-injected mice were anesthetized with isoflurane gas and euthanized with an overdose of pentobarbital (Apotekerforeningen) administered intraperitoneally, and they were thereafter perfused transcardially first with Ringer’s solution (0.85% NaCl, 0.03% KCl, 0.02% NaHCO_3_, brought to pH 6.9 with CO_2_) followed by PFA (4% in PB, pH 7.4). The brain was removed from the skull, postfixed (PFA, 4 °C, 2-24 hours) and stored in the cryoprotective (DMSO) solution. A freezing microtome was used to cut the brain into horizontal slices (40 µm or 50 µm thickness; 4 or 6 equally spaced series). All sections from brains used for neuroanatomical analysis were treated with the same basic immunohistochemical protocol described for 350 µm sections used in patch clamp experiments, with some modifications. Sections were incubated with the primary antibodies for 48 hours at 4 °C and treated with the secondary antibodies for 24 hours at 4°C or 4 hours in room temperature. Some slices were washed in a Tris buffer (2 × 10 min; 0.61% Tris(hydroxymethyl)aminomethane, pH 7.6) before being mounted on Superfrost slides (Thermo Scientific) from a Tris-gelatin solution (0.2% gelatin in Tris-buffer, pH 7.6). All other slices were washed in a mixture of PB and Tris-HCl (1 × 10 min, 1:1) and subsequently washed in Tris-HCl (2 × 10 min) and mounted on Superfrost slides using Tris-HCl.

Finally, the samples were coverslipped with either an entellan/toluene solution or an entellan/xylene solution. To reveal cytoarchitecture for correct brain area identification, a fluorescent NeuN or Nissl stain was used together with labelling for the other immunomarkers, or sections from an adjacent series were stained with cresyl violet (Nissl stain). Coverslipped samples were inspected with fluorescence illumination at the appropriate excitation wavelengths using an Axio Imager M1/2 microscope (Carl Zeiss), and digital images were obtained using an automated fluorescent scanner (Zeiss Axio Scan Z1) rigged with a Plan Apochromat (20x, NA 0.8) objective. The excitation/emission wavelengths were 355-415 nm / 412-438 nm (for FG), 450-488 nm / 501-527 nm (for AF488), 533-558 nm / 583-601 nm (for AF546) and 615-648 nm / 662-756 nm (for AF635). Confocal image z-stacks were acquired from brain sections collected from GAD67*^eCFP^* mice injected with a retrograde tracer in LEC. Image scans were taken of MEC, PIR, PER and cLEC using the same settings as described previously (see Slices from patch-clamp experiments) in order to visualize the native GFP signal of GAD67*^eCFP^* neurons along with retrogradely labelled neurons and SST neurons.

### Brain area identification

In all neuroanatomical tracing experiments, correct placement of tracer injections into the four different areas (MEC, PER, PIR and LEC) was carefully evaluated based on known cytoarchitectonic features (see below). Electrophysiological experiments were initiated after our neuroanatomical experiments had verified the correct surgery coordinates for interrogating pathways to LEC from MEC, PIR, PER and cLEC. In patch-clamp experiments, viral injections were considered to hit the intended target area whenever the axonal innervation patterns in LEC were consistent with the patterns obtained in our neuroanatomical tracing experiments. To ensure that our injections were placed in MEC, without unintended spread to LEC, we examined the innervation patterns in DG. In agreement with the current understanding of entorhinal innervation of DG in rodents (Steward, 1976; van Groen et al., 2003), injections targeting MEC or LEC resulted in axonal labelling in the middle one-third or outer one-third of the molecular layer of DG, respectively. Cases where the injection had clearly spread to LEC, evident from the laminar distribution of labelling in DG and labelled cell bodies in LEC, were excluded from analysis.

### Cytoarchitecture of EC

In this study, EC was divided into LEC and MEC, which in the rodent are distinguished by different cytoarchitectural profiles, hippocampal and extra-hippocampal connectivity patterns as well as functional properties (Nilssen et al., 2019). In short, the mouse EC is a periallocortical region made up of six layers, which includes two cell-sparse layers (I and IV) (van Groen, 2001). Across the entire EC, lamina dissecans (layer IV) separates deep layers (V-VI) from superficial layers (I -III). The laminar structure is more developed in MEC than in LEC, whereby MEC has a more regular distribution of neurons and a more distinct lamina dissecans. Neurons in the deep layers of MEC are arranged in columns, while layer II features a band of large neurons that are evenly distributed. In LEC, deep layers lack a clear columnar organization, and layer II is generally composed of two sublayers. Layer IIa is located superficial (closer to layer I) to layer IIb and contains a population of large-sized neurons that are irregularly distributed and cluster into islands. At more dorsal locations, layer IIa is separated from the smaller neurons of layer IIb by a cell-free zone. Layer IIb is made up of a densely packed population of neurons, which distinguishes this layer from the loosely arranged zone of scattered neurons in layer III. In LEC of rodents, reelin-expressing neurons distribute in layer IIa whereas calbindin-expressing neurons are found in layer IIb (Leitner et al., 2016; Nilssen et al., 2018; Doan et al., 2019; Ohara et al., 2019). Most neurons in layer III neither express reelin nor calbindin.

### Cytoarchitecture of PIR and PER

PER is a cortical area whose location in the brain is closely associated with the rhinal fissure. I ts ventral border is with LEC. A notable distinguishing feature between LEC and PER is the lamina dissecans, which disappears when transitioning from LEC to PER. Furthermore, superficial layers of PER are evenly packed with small neurons, which are replaced in layer IIa of LEC by large reelin-expressing neurons (Cappaert et al., 2015). PIR is characterized as a three-layered allocortex, consisting of an outer molecular layer (layer I) and two subjacent cellular layers. Moreover, PIR overlays the endopirifom nucleus. The three-layered appearance of PIR distinguishes it from surrounding cortical and subcortical structures. In the present study, we did not attempt to further subdivide PIR, although two main divisions (anterior PIR and posterior PIR) are often identified based on heterogeneity in cytoarchitecture and connectivity (Martinez-Garcia et al., 2012).

### Data analysis

#### Electrophysiology

All electrophysiology data analyses were performed off-line using custom-written scripts in Matlab (MathWorks). In voltage-clamp recordings, laser-evoked inward currents recorded near the reversal potential for inhibitory currents (≈-50 mV) were considered excitatory, whereas laser-evoked outward currents recorded near the reversal potential for excitatory currents (≈0 mV) were considered inhibitory. In current clamp recordings, laser-evoked excitatory responses (EPSPs) were depolarizing deflections that were observed at voltages positive to the chloride equilibrium potential (E_Cl−_ ≈ −69 mV for our solutions). Laser-evoked inhibitory responses (IPSPs) were hyperpolarizing membrane potential deflections. In cases where blockers of glutamatergic/GABAergic synaptic activity were used, the sensitivity of laser-evoked responses to the relevant pharmacological agents was used to aid classification of events as excitatory glutamatergic or inhibitory GABAergic.

In all experiments, the photostimulation protocol was repeated multiple times and the acquired traces were used to produce an average membrane current trace (voltage clamp) or membrane voltage trace (current clamp). The average trace was used to classify laser-evoked events and extract relevant features (amplitude/charge/halfwidth) of the synaptic event. In some cases (Fig. 5), the amplitude and halfwidth of the synaptic events were also calculated from the individual trials. Events with amplitudes exceeding 10 standard deviations (Z-score = ± 10; Nilssen et al. 2018) of the baseline and occurring within a 200 ms time window after laser stimulation were classified as true laser-evoked responses (EPSC or IPSC / EPSP or IPSP). Due to the noisy background resulting from concomitant action potential firing, possible laser-evoked postsynaptic potentials in the experiments forming the basis for Fig. 3 were only assessed by visual inspection. In these experiments, depolarizing/hyperpolarizing deflections that were observed in multiple trials (> 10 times) and occurred shortly after laser stimulation (<200 ms) were considered laser-evoked postsynaptic potentials. Baseline was defined as the average membrane current or voltage of a 100 ms time interval prior to laser stimulation onset. The amplitude of the synaptic event was defined as the peak value of the evoked response minus the baseline value. The response halfwidth was taken as the time difference between the rise and fall phases of the synaptic event measured at 50% amplitude. When calculating amplitude values produced by photostimulation of a given cortical layer (Fig. 2), the amplitudes of all laser-evoked responses in a particular layer were averaged. Values were then normalized by dividing this value by the maximum value of the dataset. To compare response strength across neuron groups, the synaptic charge transfer was measured. This was estimated by calculating the integral of the synaptic current (i.e., the area under the membrane current curve from the onset of laser stimulation until the membrane current returned to baseline). Membrane charge values were calculated for each photostimulation spot that produced a laser-evoked response, and for each responding neuron these values were averaged to yield a representative EPSC/IPSC charge transfer value. For non-responding neurons, i.e., neurons without laser-evoked responses, the membrane charge transfer was defined to be 0 pC.

To quantify the change in the number of emitted action potentials caused by photostimulation (Fig. 3), each recording was partitioned into 25 ms bins and for each bin the average number of action potentials across all repeats of the stimulation protocol was calculated. The relative deviation in firing activity was then calculated as the percentage of action potentials of the laser condition (during photostimulation) relative to the average number of action potentials in a 650 ms time window without photostimulation. To assess the impact of MEC photostimulation on glutamate uncaging-evoked events, the relative deviation in EPSP amplitude was calculated as the difference in EPSP amplitude (in mV) between the synaptic event evoked by combining uncaging and MEC photostimulation (375 nm + 594 nm stimulation) and the average EPSP amplitude of the two control uncaging stimulations (375 nm stimulation, without MEC photostimulation).

### Classification of recorded neurons

During patch-clamp recordings, neurons were initially categorized as belonging to either layer IIa (PN_IIa_), I Ib (PN_IIb_) or III (PN_III_) based on the location of their cell bodies relative to the cytoarchitectural profile of LEC. A final classification was performed *post hoc* by visual inspection of confocal image stacks containing biocytin-filled patch-clamp recorded neurons where the location of these neurons relative to the lamination of NeuN/Nissl cytoarchitecture and/or the distribution of reelin and calbindin was assessed. During slice recordings of GAD67^eCFP^ tissue, GABAergic interneurons were identified based on GFP fluorescence.

### Counting of retrogradely labelled neurons

To quantify the amount and nature of long-range GABAergic projections to LEC, retrogradely labelled neurons (FG/tdTomato+ neurons) within MEC, PER, cLEC and PIR were manually counted in Neurolucida 360 (MBF Bioscience) using confocal image stacks. To determine the layer distribution of neurons within MEC that had been retrogradely labelled from LEC, scanned images of MEC were inspected and tagged neurons were counted manually using Adobe Illustrator (Adobe Inc.) and assigned to the appropriate MEC layers as revealed by NeuN cytoarchitecture. All images selected for illustration purposes were imported from Zen blue/black software (Zeiss) into Adobe Photoshop (Adobe Inc.) where the contrast and brightness were adjusted.

### Counting of rabies-infected neurons

To map monosynaptic inputs to SST^+^ neurons in SST*^Cre^* mice, virus-expressing neurons were manually counted using Neurolucida coupled to a fluorescent microscope (Zeiss Axio Imager M1). One series of sections from each brain was used. Neurons expressing the rabies virus payload were distinguished from neurons expressing the helper virus payload. Neurons that contained markers from both the rabies virus and the helper virus were classified as starter cells, while neurons that expressed only the rabies virus were considered presynaptic neurons to starter cells (monosynaptic input neurons).

### Statistics

Statistical analyses were carried out in Matlab (MathWorks). Non-parametric statistical hypothesis tests were performed, as all data sets were either small or non-normally distributed as determined by the one-sample Kolmogorov-Smirnov test. All tests were two-tailed and data were considered significant at p < 0.05. The Wilcoxon signed-rank test was used on dependent samples. For independent samples, the Kruskal-Wallis test or the Wilcoxon rank sum test was applied using the multcompare function in Matlab. Bonferroni corrections were used when performing multiple comparisons.

## Supporting information

Supplementary Figure 12. Convergence of cortical inputs to principal neurons in layer IIa

Supplementary Figure 1. Dorsoventral extent of FG injection into LEC

Supplementary Figure. 2 Specificity of MEC injections.

Supplementary Figure 3. SST+ neurons project from MEC to LEC, but only very sparsely in the opposite direction, from LEC to MEC

Supplementary Figure 4. Medial entorhinal PV+ neurons and VIP+ neurons do not innervate LEC.

Supplementary Figure 5. Monosynaptic input tracing to MEC SST+ neurons

Supplementary Figure 6. Histology of representative patch-clamp recorded brain slices

Supplementary Figure 7. Layer-dependent distribution of excitatory and inhibitory synaptic inputs

Supplementary Figure 8. Pharmacological scrutiny of laser-evoked outward inhibitory currents

Supplementary Figure 9. Synaptic inputs from MEC to GABAergic neurons in LEC

Supplementary Figure 10. Synaptic inputs from MEC give rise to different responses in principal neurons in layer IIa compared with principal neurons i

Supplementary Figure 11. Anterograde tracer injections into PER, LEC and PIR

## Acknowledgements

We thank Grethe M. Olsen for technical assistance. We are also grateful to Anne Nagelhus and Øyvind A. Høydal for discussion and comments on the manuscript. This work was supported by the Research Council of Norway (research grant 227769, the Centre of Excellence scheme - Centre for Neural Computation grant 223262, and the National Infrastructure scheme - NORBRAIN grants 197467 and 295721) and the Kavli Foundation.

## Additional information

## Author contributions

Conceptualization, E.S.N., T.P.D. and M.P.W.; Investigation, E.S.N., B.J. and T.P.D.; Formal analysis, E.S.N., B.J., T.P.D. and P.J.B.G.; Visualization, E.S.N.; Software, P.J.B.G.; Writing – original draft, E.S.N.; Writing – review and editing, E.S.N., B.J., T.P.D., P.J.B.G. and M.P.W.; Supervision, M.P.W.; Funding acquisition, M.P.W.

## Funding

The Research Council of Norway (227769)

The Research Council of Norway (223262)

The Research Council of Norway (197467)

The Research Council of Norway (295721)

The Kavli Foundation

## Notes

### Competing Interest Statement

The authors have declared no competing interest.

### Summary of Updates

We have updated the manuscript based on input from two reviewers for eLife. We added supplementary material to substantiate the selectivity of the tracing experiments used for both the anatomical data as well as the in vitro electrophysiology experiments. The main finding and conclusions of the paper have not been altered however.

