## Supplementary figures and images for "Long-range inhibitory axons from medial entorhinal cortex target lateral entorhinal neurons projecting to the hippocampal formation"

### Supplementary Figure 1. Dorsoventral extent of FG injection into LEC

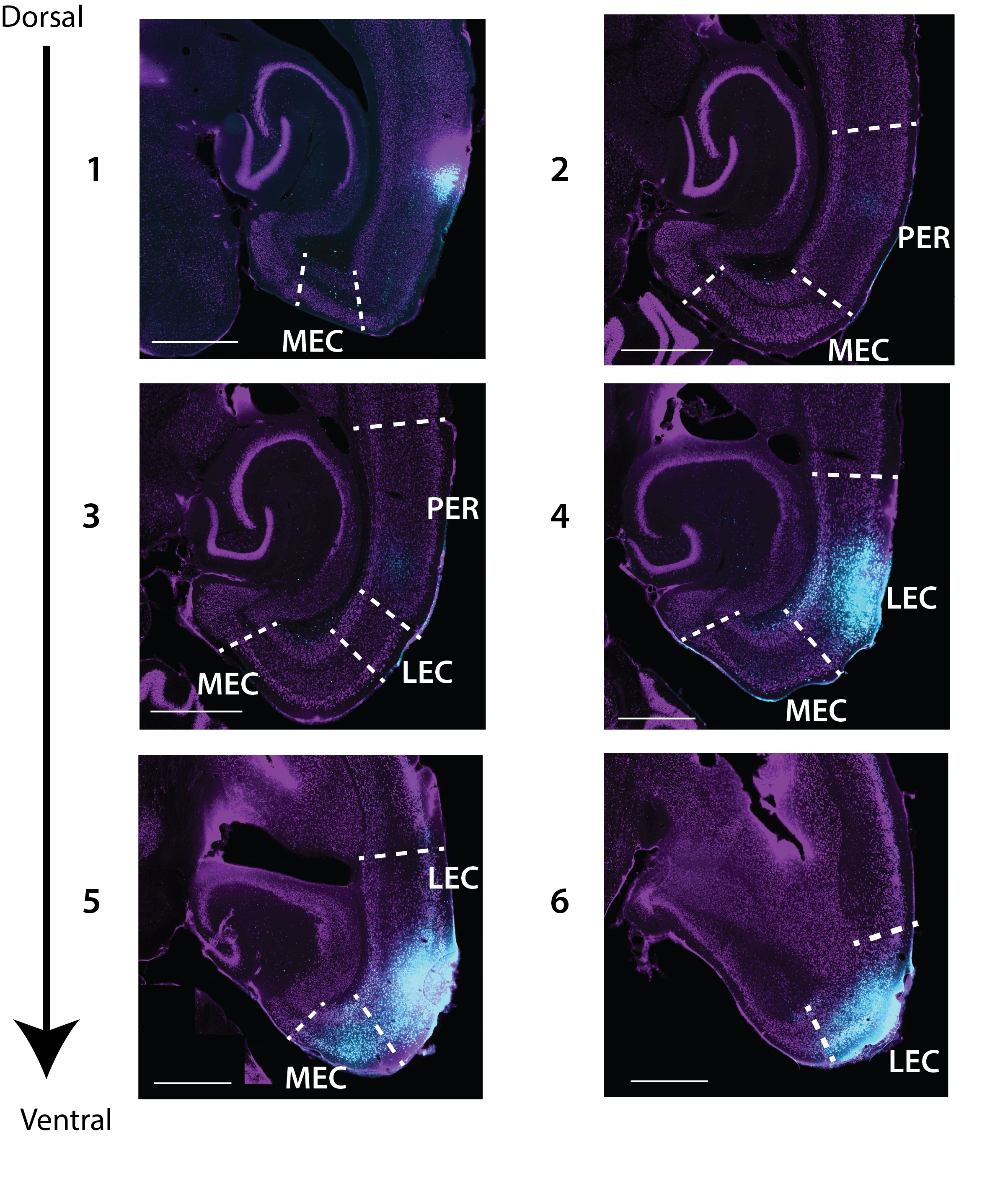

### Supplementary Figure 3. SST+ neurons project from MEC to LEC, but only very sparsely in the opposite direction, from LEC to MEC

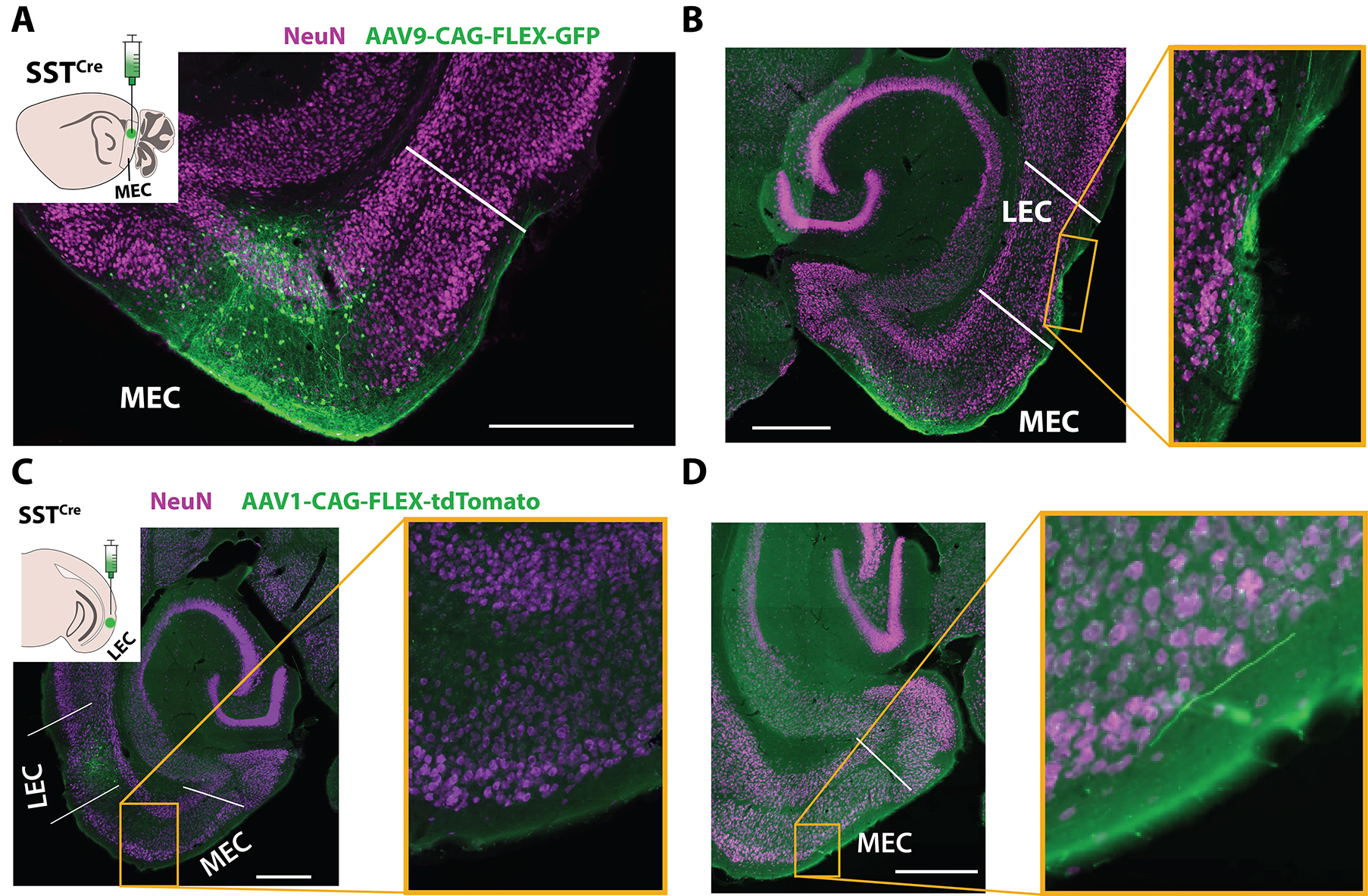

### Supplementary Figure 4. Medial entorhinal PV+ neurons and VIP+ neurons do not innervate LEC.

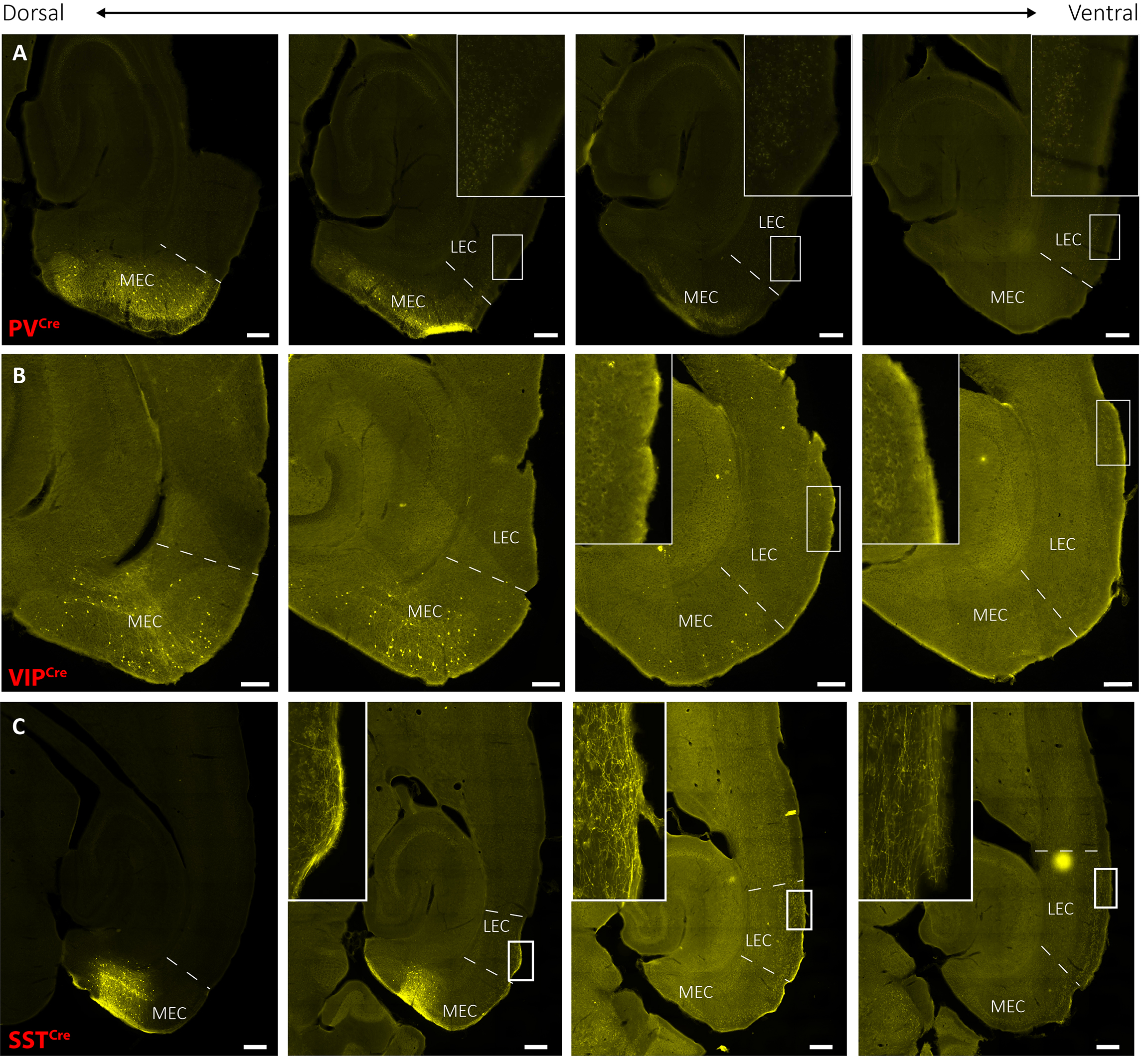

### Supplementary Figure 5. Monosynaptic input tracing to MEC SST+ neurons

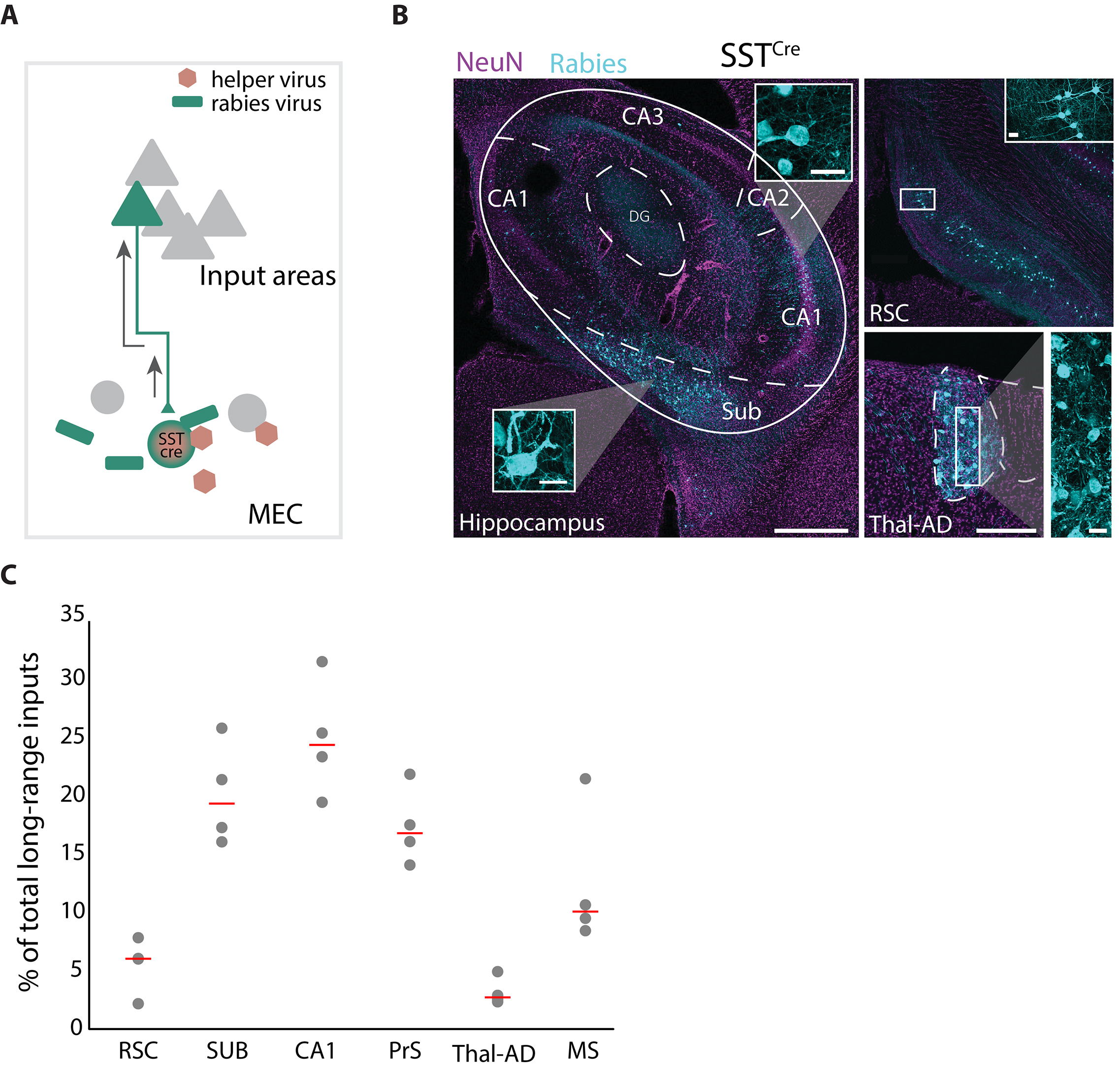

### Supplementary Figure 6. Histology of representative patch-clamp recorded brain slices

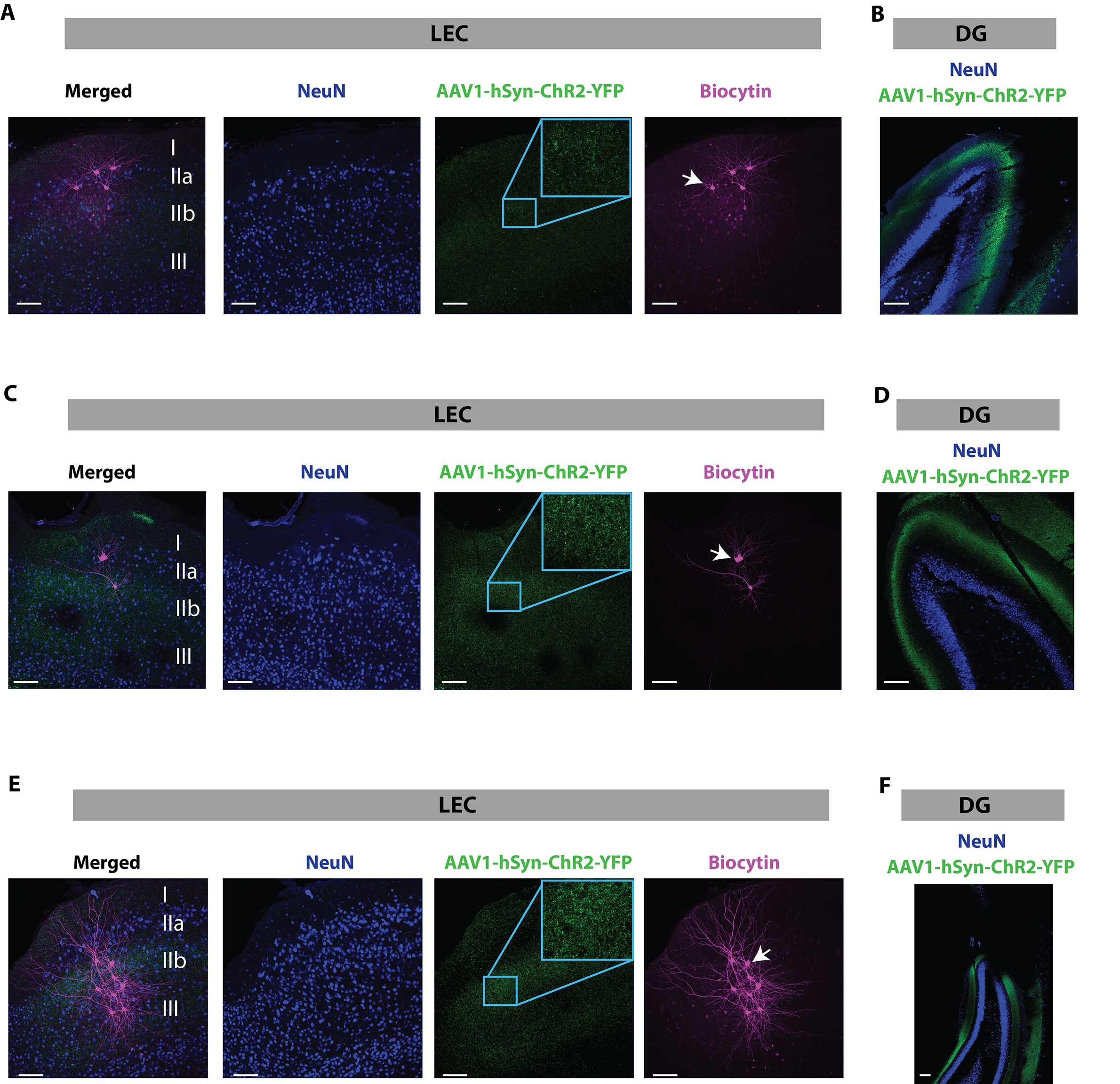

### Supplementary Figure 7. Layer-dependent distribution of excitatory and inhibitory synaptic inputs

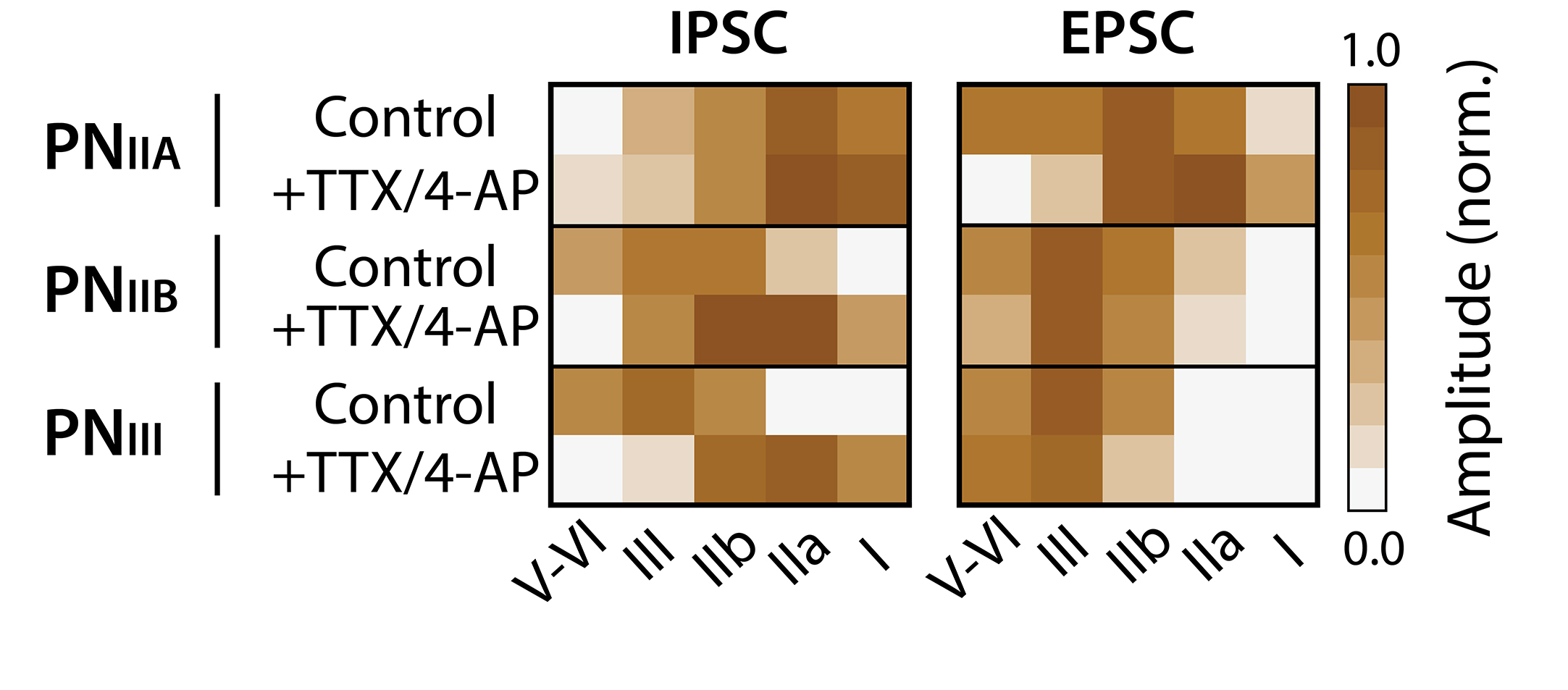

### Supplementary Figure 8. Pharmacological scrutiny of laser-evoked outward inhibitory currents

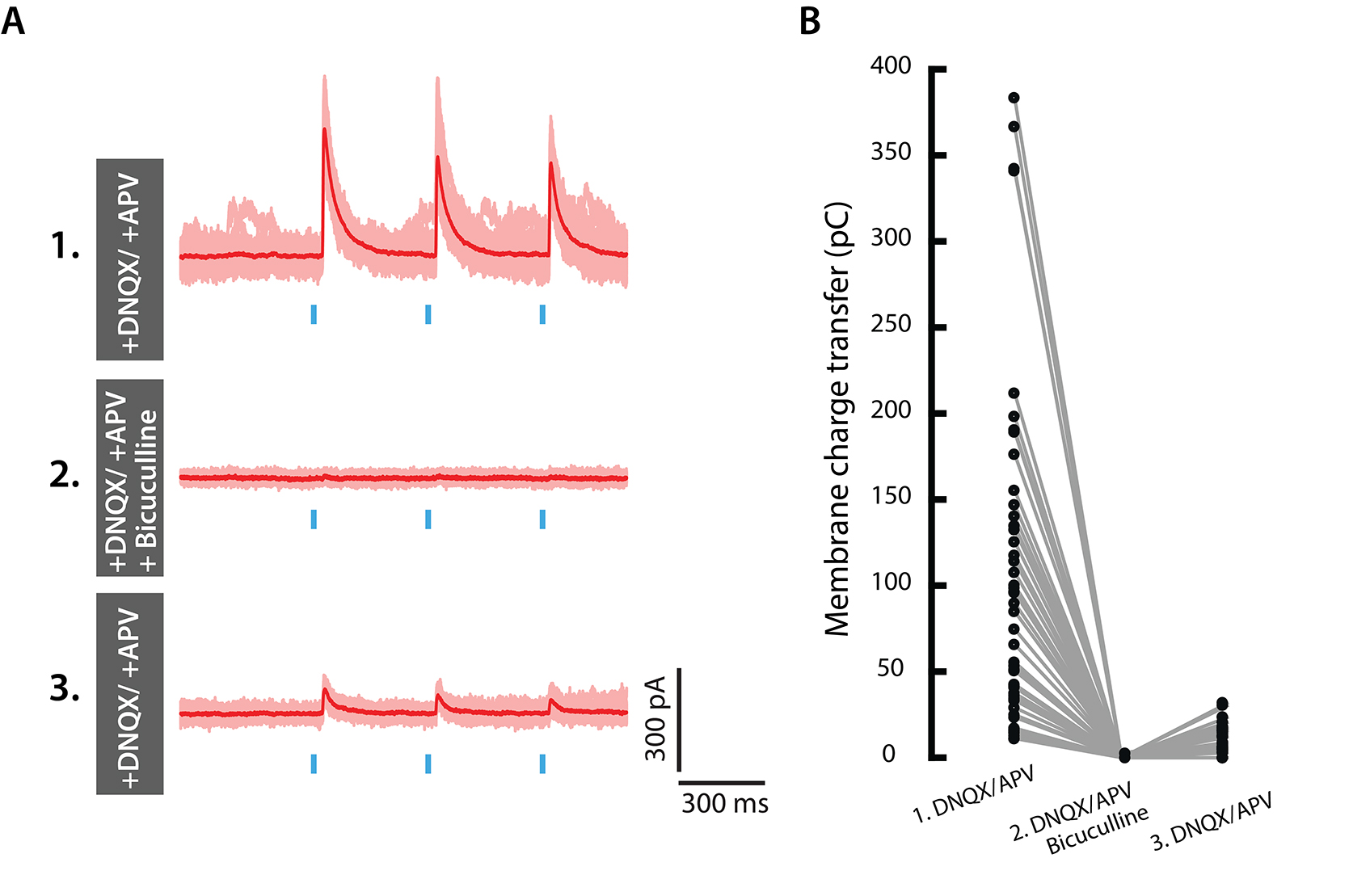

### Supplementary Figure 9. Synaptic inputs from MEC to GABAergic neurons in LEC

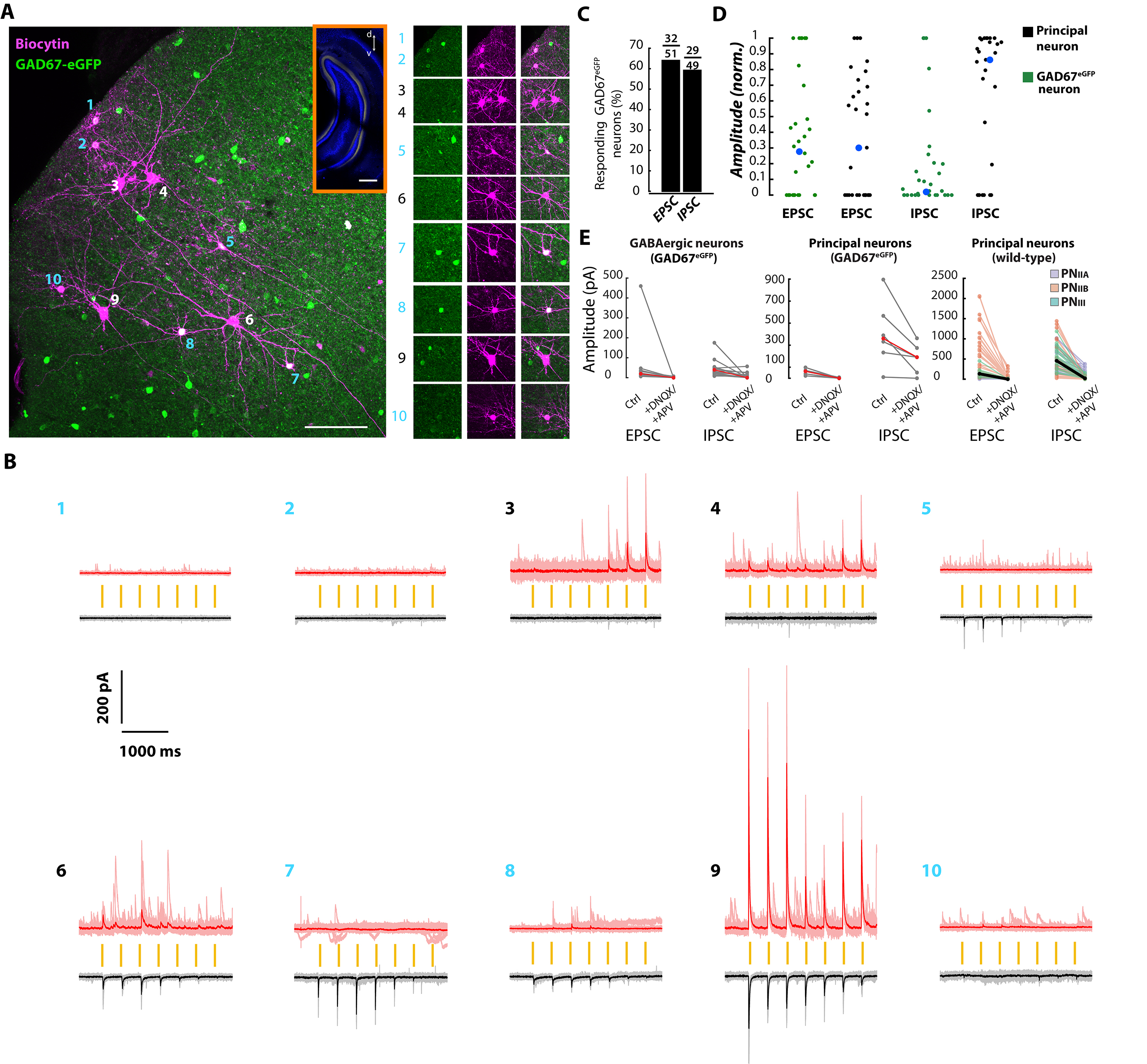

### Supplementary Figure 10. Synaptic inputs from MEC give rise to different responses in principal neurons in layer IIa compared with principal neurons i

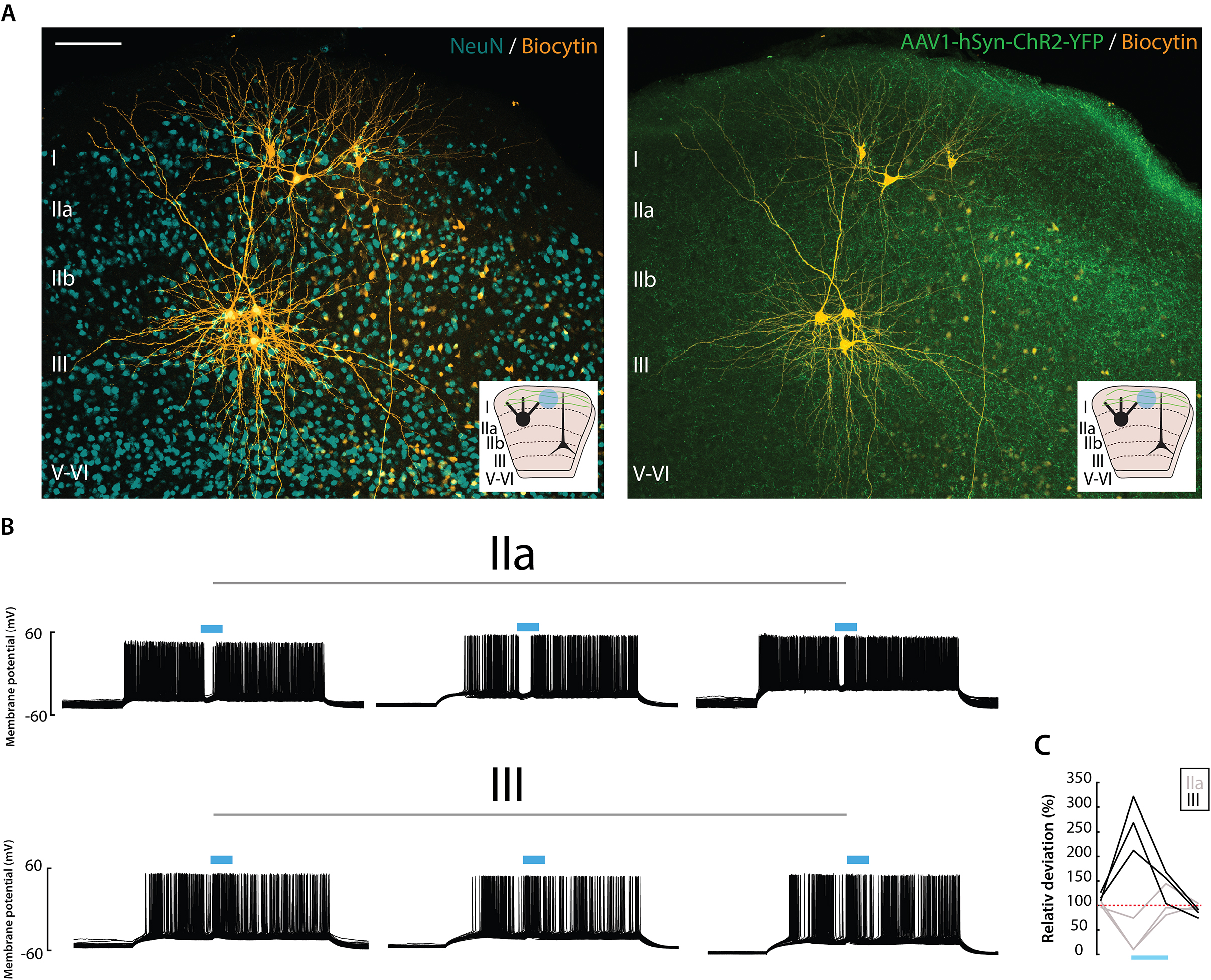

### Supplementary Figure 11. Anterograde tracer injections into PER, LEC and PIR

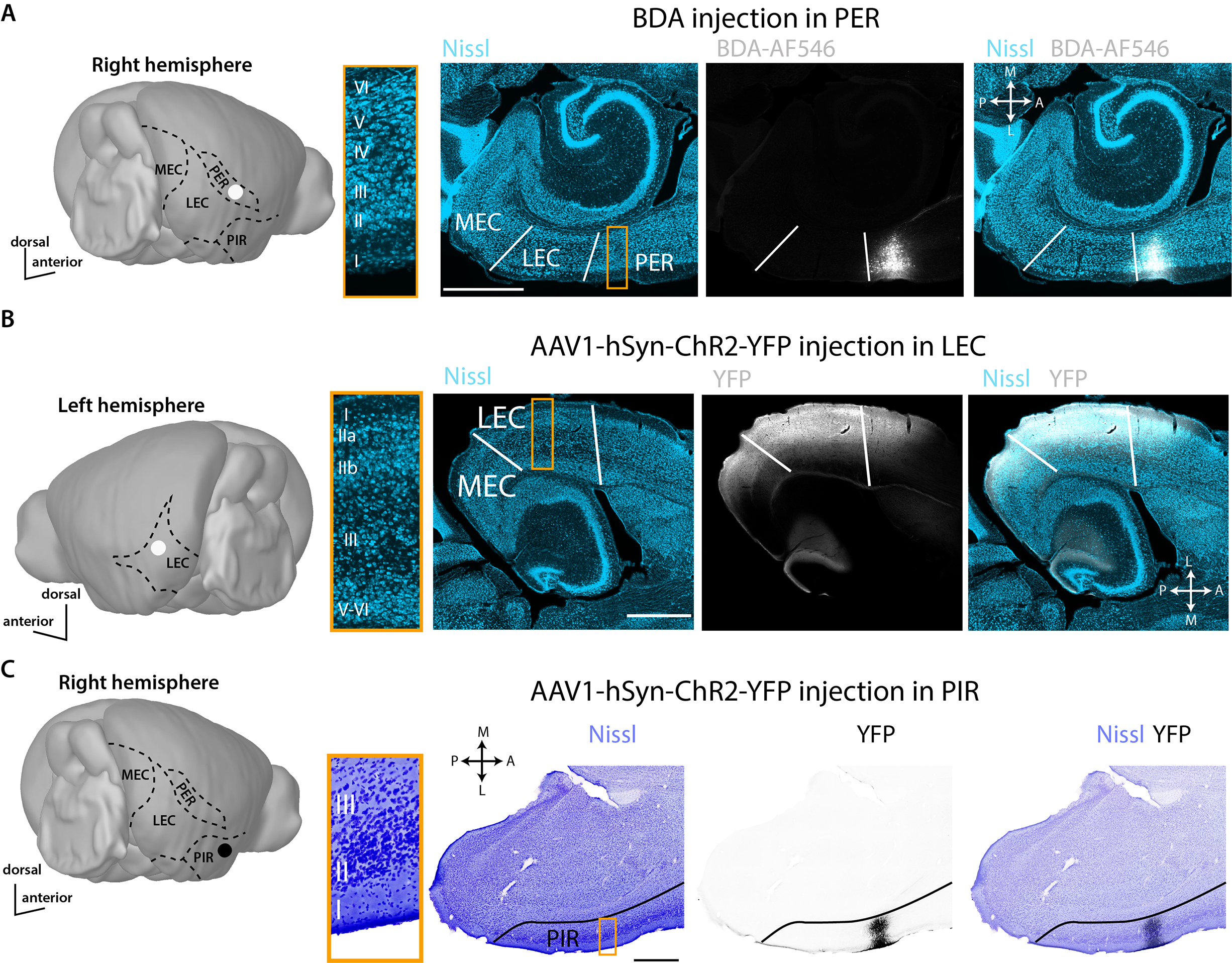

### Supplementary Figure 12. Convergence of cortical inputs to principal neurons in layer IIa

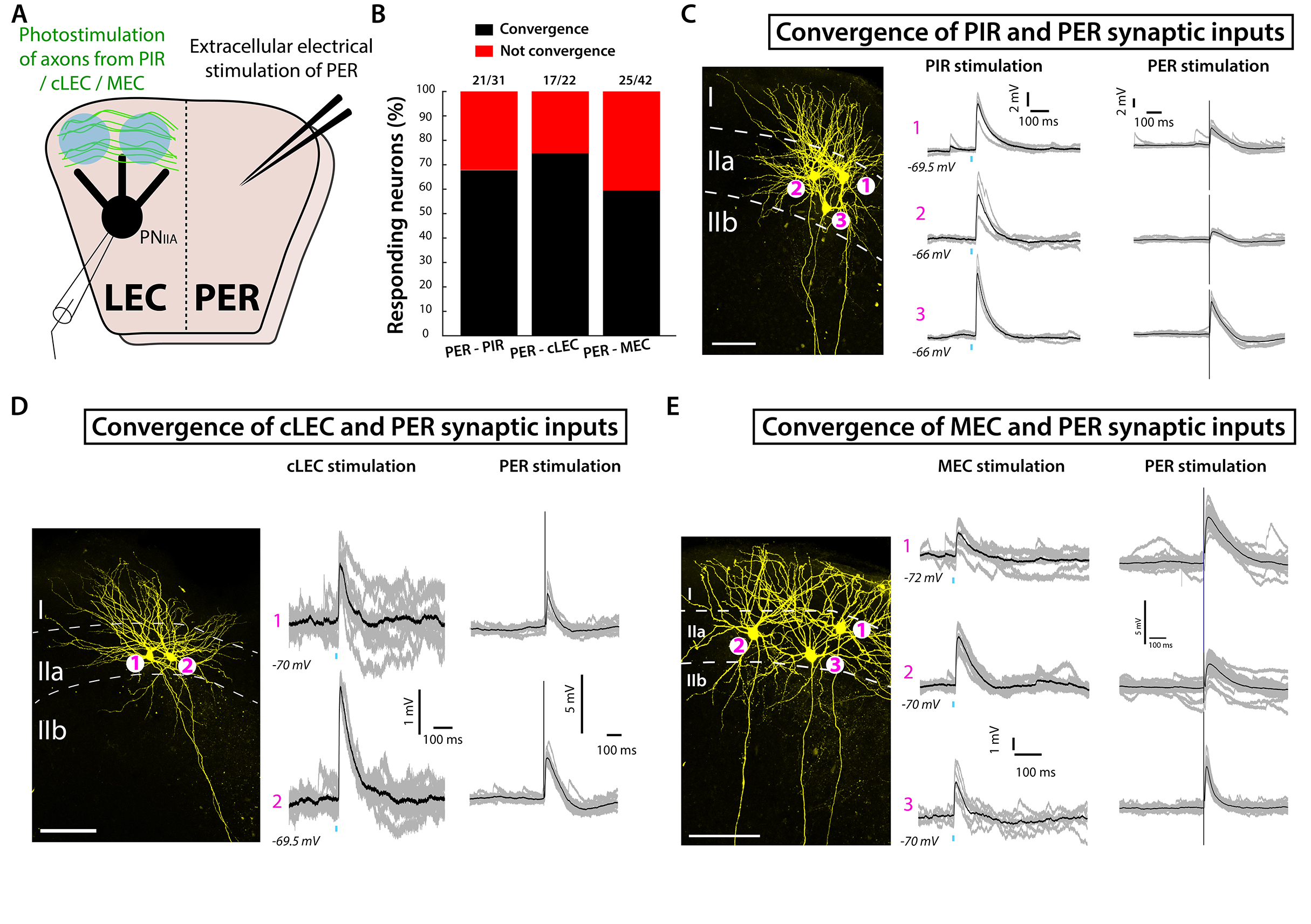

### Supplementary Figure. 2 Specificity of MEC injections.

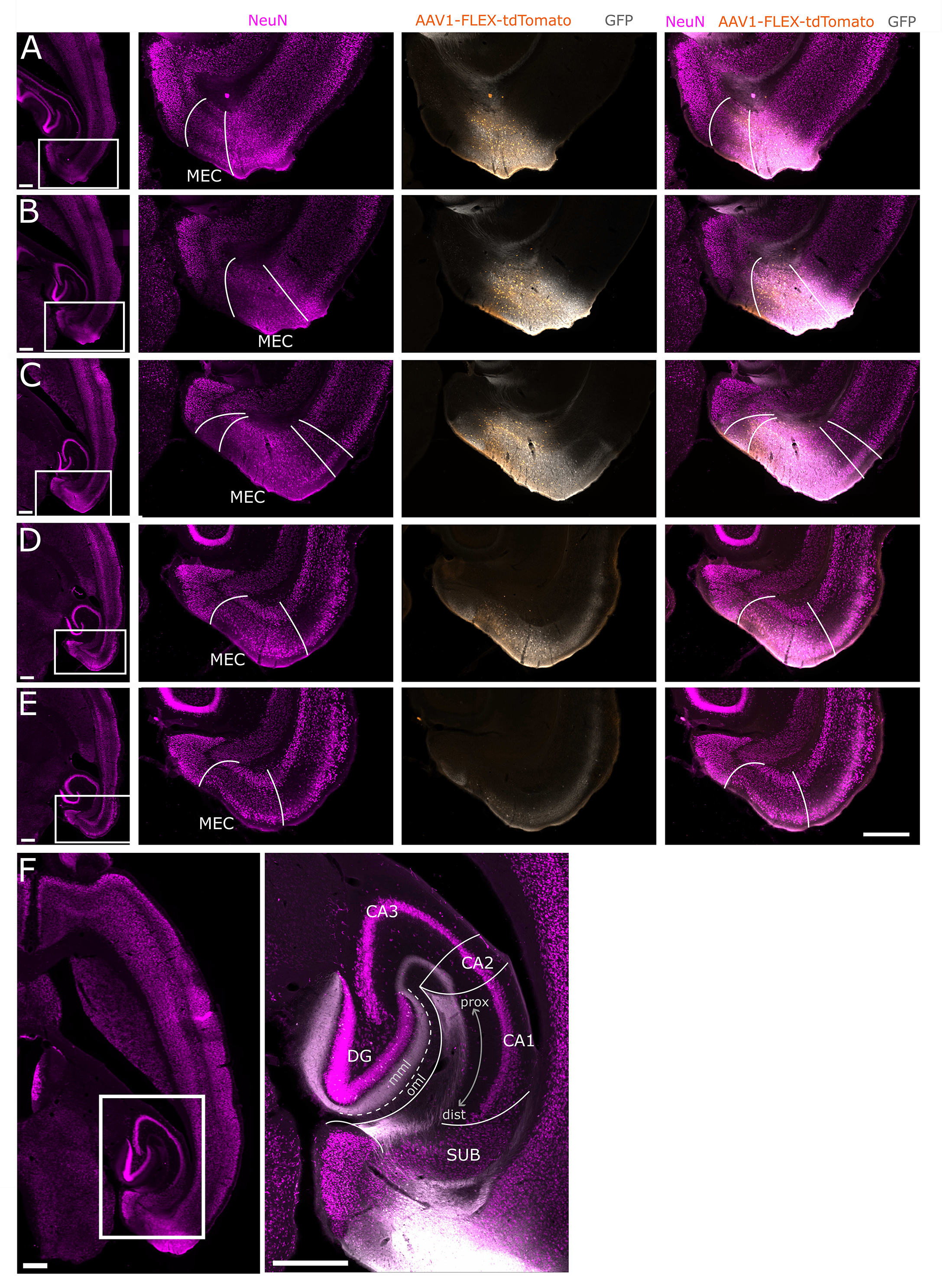
